# SORBET: Automated cell-neighborhood analysis of spatial transcriptomics or proteomics for interpretable sample classification via GNN

**DOI:** 10.1101/2023.12.30.573739

**Authors:** Shay Shimonov, Joseph M Cunningham, Ronen Talmon, Lilach Aizenbud, Shruti J Desai, David Rimm, Kurt Schalper, Harriet Kluger, Yuval Kluger

**Affiliations:** Applied Mathematics Program, Yale University, New Haven, CT, USA; Viterbi Faculty of Electrical Engineering, Technion - Israel Institute of Technology, Haifa, Israel; Department of Medicine, Yale School of Medicine, New Haven, CT, USA; Yale Cancer Center, Yale School of Medicine, New Haven, CT, USA; Yale Center for Immuno-Oncology, Yale School of Medicine, New Haven, CT, USA; Department of Pathology, Yale School of Medicine, New Haven, CT, USA

## Abstract

Spatial cellular profiling technologies have revolutionized our understanding of complex biological processes, from development and disease progression to immunity and aging. Despite their promise, integrating spatial information with multiplexed molecular data to accurately predict phenotypes poses significant challenges, especially in clinical settings. Here, we present SORBET, a geometric deep learning framework that directly analyzes complete spatial profiling data, eliminating the need to compress complete cell profiles into a limited set of annotations, such as cell types. SORBET models tissues as graphs of adjacent cells and applies graph convolutional networks to infer emergent phenotypes, such as responses to immunotherapy. The model leverages a novel data augmentation technique to ensure robust predictions, complemented by tailored interpretability analyses to identify the molecular and spatial patterns underlying the model’s phenotype inferences. We apply our method to a CosMx spatial transcriptomics dataset of pre-treatment metastatic melanoma samples annotated with response to immunotherapy; we show that spatial information significantly improves clinical endpoint, or phenotype, prediction and identifies important biological patterns. To our knowledge, SORBET is the first example of phenotype prediction on spatial transcriptomics data. We further validated our method using two spatial proteomics datasets, Imaging Mass Cytometry (IMC) and Co-detection by indexing (CODEX), obtained from Non-Small Cell Lung Cancer and Colorectal Cancer samples, respectively. SORBET demonstrates superior accuracy in phenotype prediction over leading spatial and non-spatial methods across various datasets of different observed phenotypes and technologies. SORBET sets a new benchmark for predictive analysis in spatial omics, promising to advance personalized medicine through refined patient treatment stratification, grounded in molecular and spatial tissue profiling.

## 1 Introduction

Recent advances in technologies for spatially resolved molecular profiling of cells (“spatial omics”) facilitate the simultaneous capture of both the molecular profiles of cells and their spatial organization [1–6]. The spatial organization of cells into tissues plays a crucial role in many biological processes, from embryonic patterning [7] to the progression of cancer [8]. When applied across large multi-tissue datasets, analyses that link spatial omics data to phenotypes may identify spatial expression patterns that reveal underlying biological mechanisms that differentiate the observed phenotype. However, previous studies have typically been limited to a handful of tissue samples. Moreover, modeling these data is challenging due to their heterogeneity and high dimensionality [9]. Understanding these phenotype-dependent patterns would facilitate both interrogation of molecular patterns and clinical decision making.

Graph Neural Network (GNN) models have previously been applied to analyze spatial “omics” tasks, focusing on tasks such as cell type annotation [10], cell niche clustering [11, 12], and cell-cell interaction [13]. Recently, GNN models have been extended to predict phenotypes from spatial proteomics data [14–16]. However, these GNN-based approaches reduce the rich omics profile of a cell to merely represent its cell type. To our knowledge, methods capable of utilizing the complete ‘omics profiles of cells are not available. Moreover, no methods have previously demonstrated efficacy on higher dimensional technologies, such as spatial transcriptomics platforms.

Here, we introduce Spatial Omics fRamework for Biomarker Extraction and Tissue classification (SORBET), a geometric deep learning framework designed to infer phenotypes from spatial omics data. SORBET leverages several innovations to address the challenges associated with spatial omics datasets. These include a novel data augmentation scheme to address limited sample sizes and leverage complete cell profiles. Furthermore, we introduce novel interpretability methods to understand the phenotype-specific spatial patterns identified by our model. To the best of our knowledge, SORBET is the first framework to classify multiple patient samples profiled using spatial transcriptomics at a cellular level. These data comprise metastatic melanoma tissue samples obtained prior to immunotherapy, each annotated with their corresponding response to therapy [17, 18]. Moreover, we demonstrate that our method applies across omics modalities by considering two previously published spatial proteomics datasets (CODEX, IMC) [19, 20].

SORBET incorporates several features that enhance its utility. First, the data augmentation scheme facilitates the analysis of datasets with small sample size and high-dimensional features [21]. We anticipate that this feature will allow analysis of the rapidly emerging spatial ‘omics datasets, particularly those profiled using spatial transcriptomics technologies. Second, our method utilizes the complete transcriptomics or proteomics profiles of cells, which preserves important functional information. In comparisons, existing models [14, 16] typically compress complete cell profiles into a small number of cell types. Third, we provide multiple interpretability perspectives for linking input cell profiles, cell niches, and tissue sample regions. By linking information across multiple scales, SORBET enables high accuracy classification of phenotypic groups.

In this paper, we introduce SORBET, a computational tool designed to enhance the analysis of spatial transcriptomics and proteomics data. SORBET stands out for its efficiency, accuracy, and the interpretability of phenotype predictions it provides for whole assays. As a Python package, SORBET is readily accessible and seamlessly integrates into existing spatial omics analysis pipelines. The development of SORBET is a step towards expanding the toolkit available to researchers in the field of biomedical research, potentially unlocking novel insights into the mechanisms underlying health and disease.

## 2 Results

### 2.1 SORBET Overview

SORBET, a supervised geometric deep learning framework, learns the relationship between spatial ‘omics data and phenotypic labels. These labels correspond to various biological properties or clinical endpoints related to the tissue samples such as response to therapy (binary variables), histologic subtype (categorical), or progression-free survival (continuous). We started by applying SORBET to classify responses to immunotherapies for pre-treatment tissue samples derived from melanoma patients (8.8). The input data for SORBET comprise three elements: phenotypic labels, the spatial coordinates of cells, and the omics profile of each cell (8.1). For clarity, we split SORBET into three steps: ‘Graph Modeling’, ‘Learning’, and ‘Reasoning.

#### 2.1.1 Graph Modeling Step

In the Graph Modeling step, tissues are modeled as graphs, with each cell corresponding to a “node” and cell-to-cell contact represented as an “edge” (**Fig. 1a**; 8.2). We chose a graphical model to simultaneously capture both cell profiles and the context of their surrounding cells. However, training supervised deep learning methods on whole tissue samples poses a challenge due to limited sample sizes. Rather than model whole tissues, we divide each sample into automatically selected subregions, or ‘subgraphs’ (center, multi-colored sample, **Fig. 1a**; 8.4). Briefly, the algorithm first identifies ‘anchor’ cells in the tissue (top left, colored cells, **Fig. 1b**). Anchor cells may be identified randomly or may incorporate prior information. Prior information, such as tumor markers in the context of cancer, may be used to guarantee a non-negligible fraction of certain cell types (*e*.*g*., cancer cells) are present in each subgraph. For the melanoma dataset, we identify anchor cells with high expression of *S100B*, a melanoma tumor marker. Next, the algorithm iteratively expands around each anchor cell to extract representative subgraphs from each tissue (right, change from black to color, **Fig. 1b**). Each subgraph is labeled with the phenotype of the sample from which it originates and is used as input to the subgraph prediction task. The subgraph extraction algorithm, a type of data augmentation, extracts multiple subgraphs from each original tissue (overlapping regions may be selected; see ‘Subgraph Extraction’ tissue sample, **Fig. 1a**). Data augmentation techniques, such as contrastive learning or partitioning images into patches, have been applied in other modeling domains to improve model performance in the context of limited sample sizes[21].

**Fig. 1.**
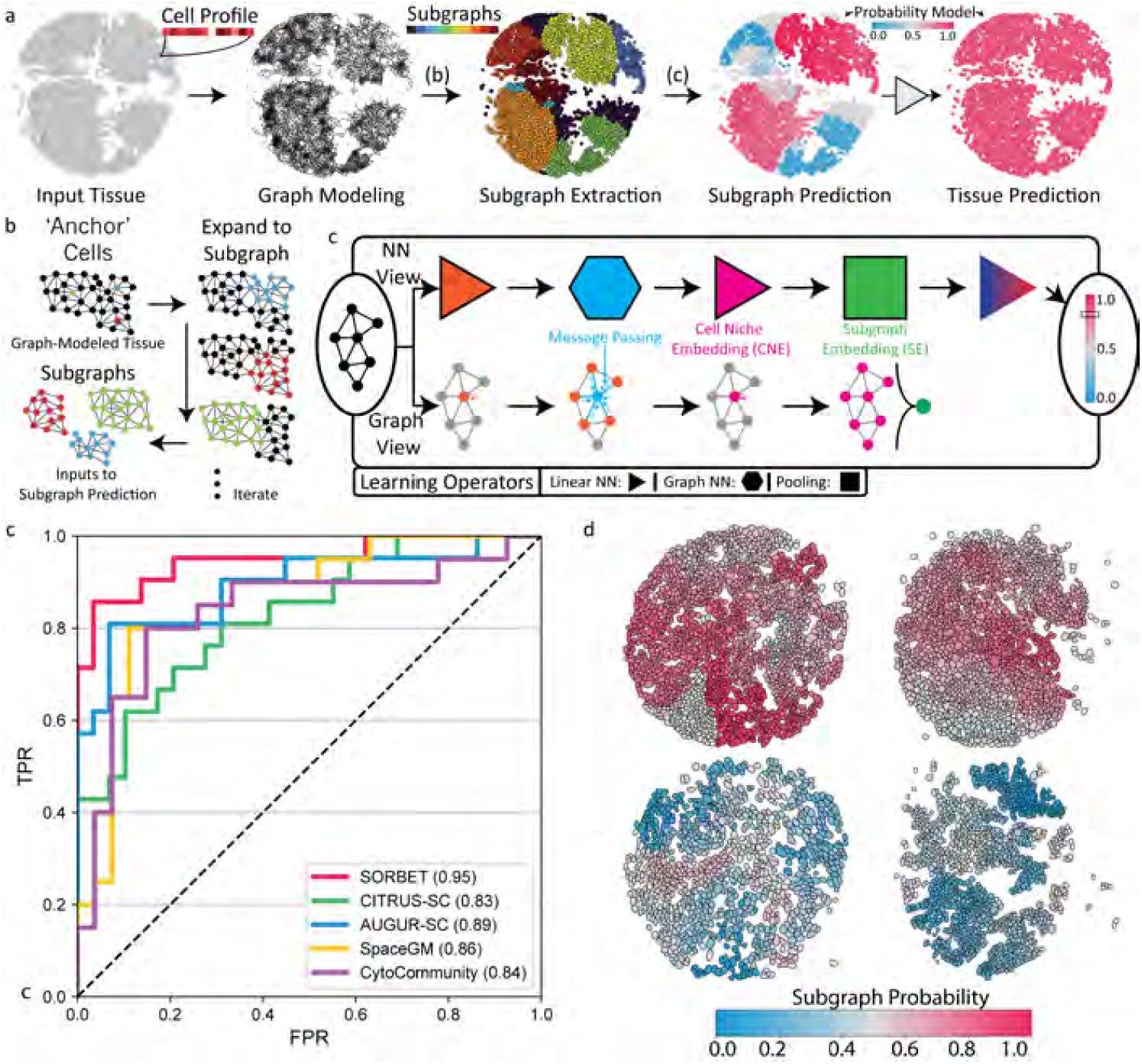
SORBET Model and Performance Evaluation Predicting Response to Immunotherapy for Pretreatment Metastatic Melanoma Tissue Samples Profiled Using CosMx (Spatial Transcriptomics) **(a)** (left) Input tissues, characterized by the spatial coordinates of cells and the omics profile of each cell, are modeled as graphs (middle left, ‘Graph Modeling’), with an edge corresponding to cell-to-cell contact. (right) Applying a data augmentation strategy, each input graph-modeled tissue is automatically split into subgraphs (‘Subgraph Extraction’, see **(b)**). A label is predicted for each extracted subgraph using a GCN-based model (middle, ‘Subgraph Prediction’, see **(c)**) and the predictions are combined into a prediction for each tissue (right, ‘Tissue Prediction’). **(b)** Subgraph extraction algorithm. ‘Anchor’ cells are identified in each tissue either randomly or with prior information (*e*.*g*., a tumor marker). Small tissue regions, or subgraphs, are iteratively extracted around each anchor cell. Each subgraph is labeled with the phenotype of the sample from which it originates and is used as an input to the ‘Subgraph Prediction’ task. **(c)** GCN-based model architecture. Subgraphs are passed through four neural network operators (‘Learning Operators’), resulting in a predicted subgraph label and associated confidence level (right). Operations are plotted using both a neural network (‘NN’) view (top) and a graphical view (bottom). A fully-connected NN (FCNN) computes an embedding of each cell’s molecular profile (NN View: triangle operator, Graph View: orange arrow to self). A representation of the cell’s niche, the CNE, is computed by convolving with the profiles of nearby cells (hexagon operator, blue arrows). A max-pool operation aggregates CNEs within each subgraph into a single subgraph representation, the subgraph embedding (SE; square operator, green node). SEs are passed through an FCNN to compute a predicted label and confidence level for each subgraph(probability scale, right).**(d)** Patient classification performance assessed on Melanoma Spatial Transcriptomics Dataset. Curves represent the mean of ten random re-initializations for five-fold cross validation. Performance is recorded in the lower-left legend. Compared against four methods: two GNN methods (SpaceGM, CytoCommunity) developed for spatial proteomics and two classical methods (AUGUR, CITRUS) developed for (non-spatial) single-cell transcriptomics. **(e)** Subgraph predictions are plotted for four tissue samples. Each cell (black border) is color-coded to reflect the estimated likelihood that the subgraph to which it is associated will respond to immunotherapy (‘Subgraph Probability’ colorbar; blue - low likelihood, red - high likelihood). Cells that appear in multiple overlapping subgraphs are colored by the mean predicted probability from all sub-graphs that they belong to. Any cells not assigned to any subgraph are set to 0.5 (grey). The top two samples belong to responders to immunotherapy; the bottom two samples belong to non responders to immunotherapy.

#### 2.1.2 Learning Step

In the Learning step, subgraphs are classified using a graph convolutional network (GCN) based architecture (**Fig 1a,c**; 8.6) [22–24]. The GCN model learns a new representation for each cell that convolves the cell’s profile and the profiles of nearby cells (blue arrows, **Fig. 1c**; 8.5). This representation, which we term cell niche embedding (‘CNE’; pink nodes, **Fig. 1c**), captures the ‘omics profiles of the cell and its local niche. Next, we use a max-pooling operation to aggregate all the cell niche embeddings within each subgraph into a single representation, which we term subgraph embedding (‘SE’; green nodes, **Fig. 1c**). By construction, the subgraph embeddings share the same space as the cell niche embeddings. Subsequently, subgraph embeddings are passed through a fully-connected neural network, which yields a prediction of each subgraph’s phenotype along with the corresponding level of confidence (middle, ‘Subgraph Predictions’, **Fig. 1a**). Finally, the predicted sample label is computed as a sum of the predicted subgraph labels weighted by their confidence (right, ‘Tissue Prediction’; **Fig. 1a**).

#### 2.1.3 Reasoning Step

In the Reasoning step, we analyze patterns at multiple biological scales inferred by SORBET using novel interpretability approaches (8.7). Specifically, we analyze the relationships between the spatial omics input data, cell niche embeddings, and subgraph embeddings. First, the relationship between cell niche embeddings and predicted phenotypes is quantified using a score we term Cell Phenotypes Similarity (CPS; **Fig. 2a, Section 2.3**). With this CPS, we highlight tissue structures that are predicted to be associated with each phenotype. We extend these analyses using cell type annotations (**Fig. 3a,b**). The CPS, when combined with cell type annotations, identifies repeated spatial, typed cell patterns across tissues correlated with predicted phenotypes. Second, we leverage sparse Canonical Correlation Analysis (sCCA) [25] to compute the relationship between the input data (orange nodes, **Fig. 4a**) and each cell niche embedding (pink nodes). This analysis identifies the most relevant individual markers or groups of markers that differentiate each phenotype. Using these marker groups, we identify the types of biological patterns, such as pathways or gene families, that the model uses to generate for accurate predictions.

**Fig. 2.**
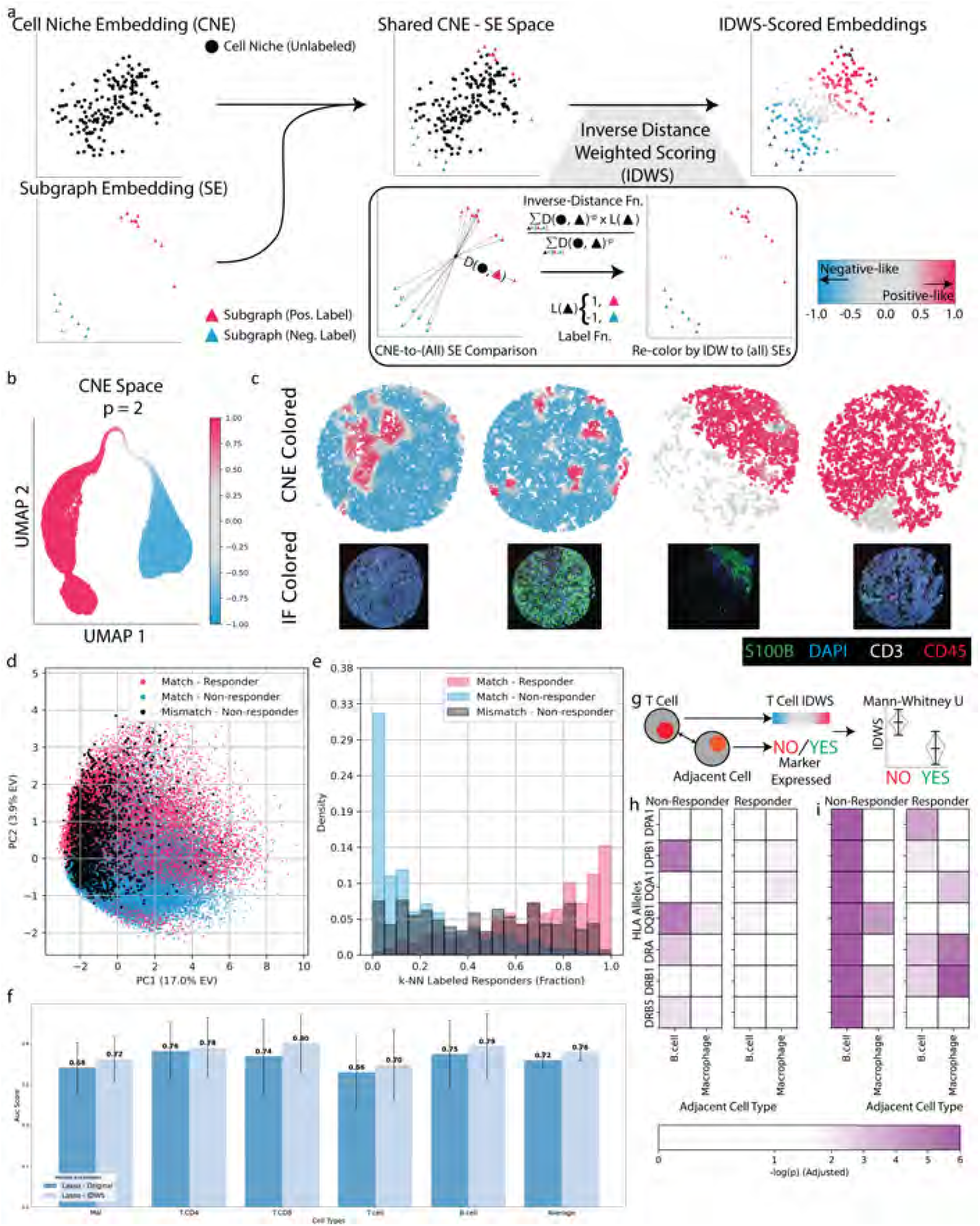
CPS analysis identifies spatial patterns of response to immunotherapy in metastatic melanoma. **(a)** Computation of Cell Phenotypes Similarity scores (CPS) from cell niche embeddings (CNE) and subgraph embeddings (SE). CNEs and SEs occupy the same embedding space. CPS characterizes cell niche similarity to subgraphs and, by extension, phenotype, ranging from −1 (non-response; blue) to 1 (response; red). **(b)**UMAP embedding of all CNEs, colored by CPS (*p* = 3). **(c)** Representative tissue sections from responders (*n* = 2; left) and non-responders (*n* = 2; right). Top: Cells colored by CPS, as shown in (b); bottom: immunohistochemistry staining of same tissue sections. **(d)** Principle component analysis of CNEs (|CPS| *>* 0.5) showing three groups: responder tissues with CPS *>* 0.5 (red), non-responder tissues with CPS *<* − 0.5 (blue), and non-responder tissues with CPS *>* 0.5 (black). **(e)** *k*-nearest neighbor analysis (*k* = 50, “k-NN”) showing fraction of neighbors from responder tissues for each group in **d. (f)** Classification of cells, grouped by cell type, by tissue of origin. Classifiers trained for either all cells (‘Original’) or for cells filtered by CPS threshold (‘CPS’, |CPS| *>* 0.5). **(g)** Schematic of T cell activation analysis comparing CPS distributions contingent on MHC expression in adjacent APCs (Mann-Whitney U test). **(h,i)** Analysis of T cell CPS distributions for **(h)** Th1 CD4^+^ T cells near B cells or macrophages or **(i)** CD8^+^ T cells near non-professional APCs (malignant cells, cancer-associated fibroblasts, or endothelial cells). log_10_(p) adjusted shown (Benjamini-Yekutieli adjustment). Color separates non-significant (white, adjusted *p >* 0.05) from significant (shades of purple, adjusted *p <* 0.05)

**Fig. 3.**
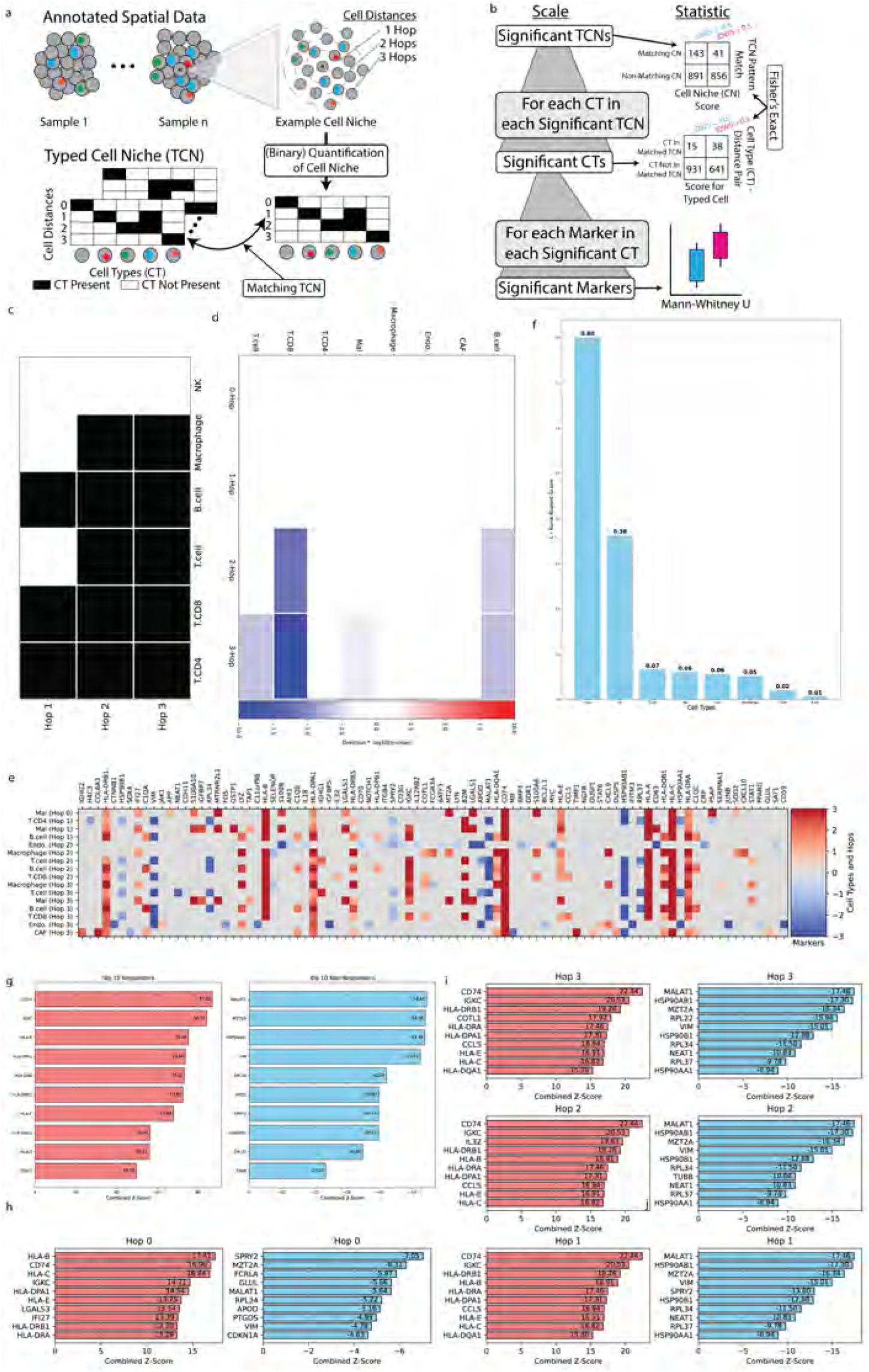
Spatial cell-type patterns distinguish immunotherapy response in metastatic melanoma. **(a)** TCN analysis framework. Three-hop neighborhoods are extracted around malignant cells (top), with distances marked by concentric rings. Each neighborhood is encoded as a binary matrix (bottom) indicating presence/absence of cell types at each distance. **(b)** Multi-scale statistical analysis cascade: Fisher’s Exact Tests identify (top) significant Typed Cell Niches (TCNs) and (middle) significant cell type-distance (C-D) pairs within TCNs; (bottom) Mann-Whitney U tests identify differentially expressed markers within significant CT-Distance pairs. **(c)-(e)** Analysis of representative non-responder-associated TCN showing **(c)** binary pattern matrix, **(d)** significance of CT-Distance pairs (− log_10_(P values)), and (e) median expression of top differentially expressed markers (*n* = 10) in each significant CT-Distance pair. **(f)** Bar plots comparing the Ranked Biased Overlap (RBO) between significant markers identified for each cell type using either the original labels or CPS scores. **(g)** Expression of the top 10 differentially expressed markers across all CT-Distance pairs and all TCNs in (left) responders and (right) non-responders. **(h,i)** Expression of the top 10 differentially expressed markers across all TCNs in **(h)** central malignant cells and **(i)** CD8+ T cells at each hop distance (1-3 hops) for (left) responders and (right) non-responders.

**Fig. 4.**
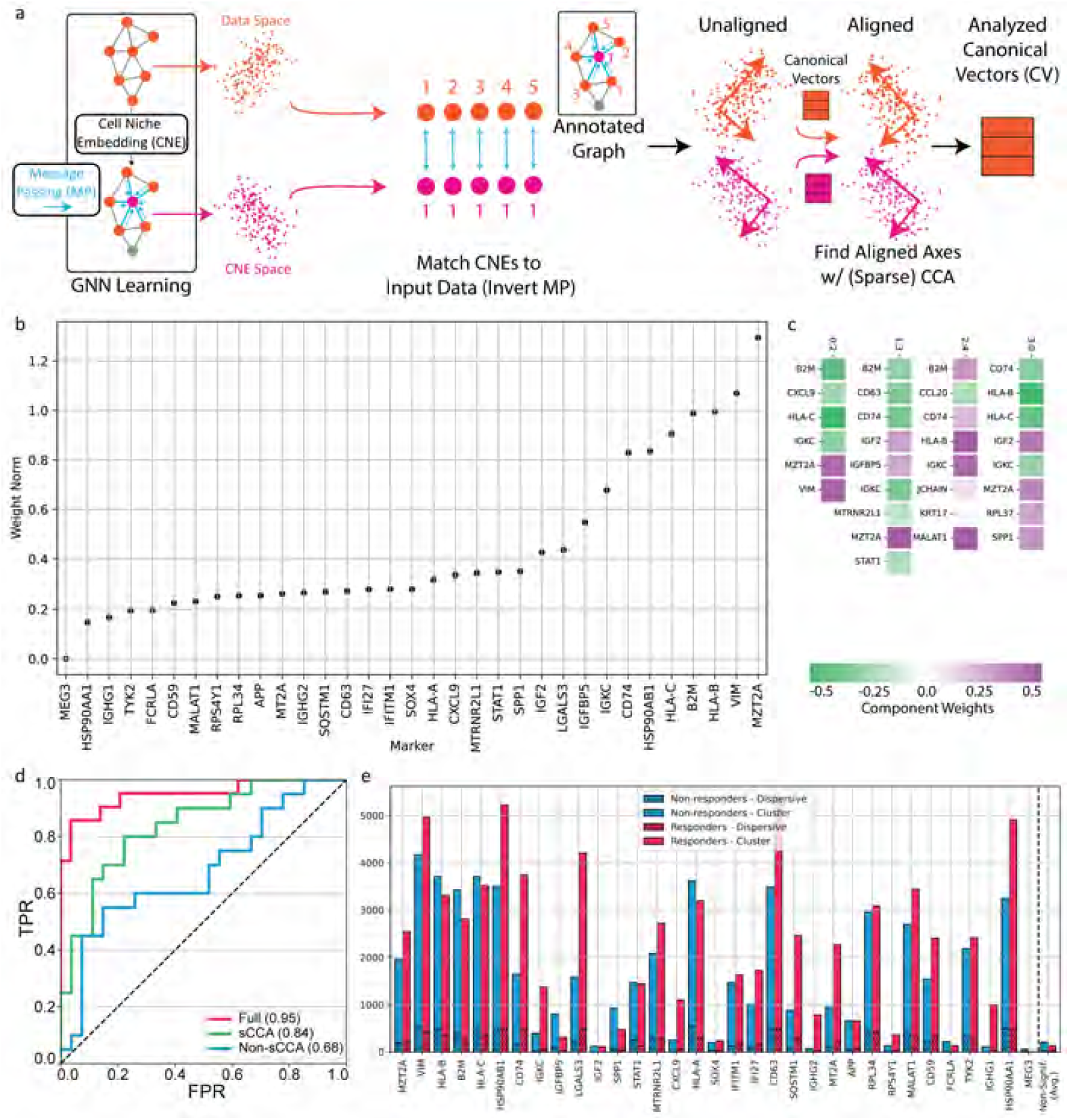
Spatial marker patterns that distinguish immunotherapy response in metastatic melanoma. **(a)**Sparse canonical correlation analysis (sCCA) framework. CNEs (pink) are computed from three-hop neighborhood transcriptional profiles (orange) through message passing. CNEs are paired with input profiles based on hop distances, enabling identification of aligned axes between input and CNE spaces. **(b)** Marker prioritization across spatial scales combining weights from five sparse components at each distance (0-3 hops). Overall contribution, or Weight Norm, reflects each marker’s importance across all spatial scales for predicting treatment response. **(c)** Representative marker programs at each graph distance, labeled as ‘Distance:Component Index’. **(d)** Classification performance comparison of SORBET using full dataset (red; same as **Figure 1c**), sCCA-identified markers only (green), or excluded markers only (blue). **(e)** Local spatial association analysis of sCCA-identified genes. Number of significant cells with dispersive (Local Moran’s I *<* 0; dots) or clustering (I *>* 0; solid color) statistics for tissues from responders (red) or non-responders (blue). P values adjusted using Benjamini-Yekutieli method. (right of dashed line) Significant cells for all other markers (average).

### 2.2 SORBET Accurately Classifies Immunotherapy Resistance Versus Sensitivity in Tumors From Spatial Transcriptomics Profiling Data

To evaluate phenotype classification accuracy on spatial transcriptomics data, we analyzed a dataset comprising 47 patient-derived melanoma samples profiled using the CosMX Spatial Molecular Imager [6] platform (**Table ??**; 8.8.1). Tumor samples were prepared in a tissue microarray format, as described [17, 18]. Each patient received one of two immunotherapy types (Ipilimumab and Nivolumab combination therapy, or anti-PD1 monotherapy (Pembrolizumab or Nivolumab)). Responders (n = 20) were defined as patients demonstrating a partial or complete response to therapy, as determined by comprehensive body scans using radiologic response criteria (RECIST), while non-responders were those with stable or progressive disease [17, 18].

Model accuracy was assessed using a *k*-fold cross-validation procedure (*k* = 5; 8.6). The cross-validation procedure is stratified by two factors. First, subgraphs derived from the same tissue are sorted into the same cross-validation fold. This strategy prevents data leakage between data used for training and testing models. Second, positively and negatively labeled samples are balanced across folds. This choice is required due to the small dataset size. For each dataset, cross-validation is repeated ten times. Each repeat is a new random split of the data. The mean and standard deviation of the receiver-operator characteristic (ROC) curve and Area Under the ROC (AUROC) curve are reported (8.6).

For comparison, we consider four, previously reported baseline methods. Two recently published methods, CytoCommunity [16] and SPACE-GM [14], are GNN-based methods that leverage spatial information and processed cell profiles (*e*.*g*. cell types). These methods were previously applied to a published spatial proteomics dataset (“CRC” dataset; analyzed in **Sec. 2.7**) [14, 16]. We extend these methods to both the metastatic melanoma spatial transcriptomics task and a new spatial proteomics task (“NSCLC” dataset, **2.7**). To the best of our knowledge, all existing GNN-based techniques fail to utilize the complete omics profiles of cells; rather, these methods typically use limited representations, such as cell types. In addition, we compared our spatially-informed method to two methods, CITRUS [26] and Augur [27], that were developed for single-cell omics data without spatial information.

SORBET achieves an AUROC of 0.95 (**Fig. 1d**) outperforming both the non-spatial single-cell omics methods (Augur-SC: 0.89 (AUROC), CITRUSSC: 0.83) and the spatial GNN-based methods (CytoCommunity: 0.84, SPACE-GM: 0.86). We conjecture that the benefits of SORBET mainly result from two factors. First, SORBET employs the full omics profiles, which allows for greater model expressiveness in contrast with methods such as SPACE-GM and CytoCommunity that reduce the full omic profiles into limited cell annotations. This is crucial as the full cell omics profile defines not only the type of each cell but also its functional state. Second, SORBET utilizes a data augmentation technique tailored for spatial biological datasets.

Use of subgraphs rather than whole tissues, provides flexibility to model tissue heterogeneity and identify spatial regions within a single sample that may be classified as opposite to the tissue’s phenotype. To illustrate sample heterogeneity, we highlight the subgraph predictions of SORBET in four melanoma samples (**Figure 1e**). Here, the model prioritizes multi-cellular sub-regions associated with response (or non-response) to immunotherapy. The two samples on the top correspond to samples from patients who responded to the therapy. For these samples all subgraphs were predicted to be sensitive to immunotherapy with high probability. In contrast, most subgraphs in the two bottom samples, which were from patients who did not respond to immunotherapy, are predicted not to respond to therapy. However, in one of these samples (bottom left, **Fig. 1e**), some of the subgraphs have higher likelihood scores, implying that some parts of the tumor may have been sensitive to immunotherapy whereas other portions were associated with tumor out-growth. Nevertheless, the model predicts both of the bottom samples have a low likelihood of responding to therapy. We hypothesize that the difference in predicted response to therapy is consistent with previous observations that heterogeneity within tumors can limit response to immunotherapy [28–31].

In clinical practice, a tool like SORBET would be required to generalize to novel patient populations. Unfortunately, a true validation set is not available to assess the model’s capacity to generalize. However, during data collection, the metastatic melanoma cohort was sequenced in two separate batches. Although not independent, this experimental set-up provides an approximate validation set. We re-trained SORBET on the first batch, which consists of approximately half of the data (*n* = 27). The re-trained model achieves high predictive accuracy (AUROC = 0.82; Supplementary Figure **??**). This performance, which entails generalization from a small dataset, implies that SORBET may effectively serve as a biomarker across heterogeneous data.

### 2.3 Classification Interpretability via Identification of Spatial and Molecular Patterns

The Reasoning Step of SORBET analyzes three embedding spaces: the input data space, the inferred cell niche embedding, and the subgraph embedding spaces (8.7). These spaces represent different biological scales from molecular information of each cell in the input space to inferred representations of cell niches and sub-tissue regions. Our interpretability analysis is split into two core methods: one focuses on spatial cell patterns and the other focuses on spatial markers patterns. These two methods capture different spatial aspects of the phenotype prediction. Analysis of spatial cell patterns identifies key tissue structures at different scales. In a similar vein, analysis of spatial marker patterns detects the key marker programs and assesses the individual, spatial contributions of each marker. The analysis identifies both unique and shared patterns across multiple individuals. As opposed to typical methods, these novel GNN interpretability techniques rely solely on the learned embeddings without the need to re-train additional models or monitor the training process.

#### 2.3.1 Spatial Cell Patterns

To discern key tissue structures at the cell and cell niche levels, we devise a correlation score for each cell niche embedding with the predicted phenotype of the tissue. As demonstrated in **Fig. 1d**, the model accurately predicts the phenotype for each subgraph embedding, which represents a distinct subgraph. By choice of pooling operator (`max-pool`), the subgraph and cell niche embeddings share the same space (left and middle, **Fig. 2a**). As a result, we may extend the likelihood of each phenotype score to cell niche embeddings, by measuring the similarity of each cell niche embedding to all labeled subgraph embeddings (‘CNE-to-(All) SE Comparison’, boxed inset, **Fig. 2a**). To compute this similarity, we utilize an inverse-distance weighted score, which we term Cell Phenotypes Similarity score (CPS; 8.7). The weighting gives greater emphasis to subgraph embeddings closer in distance (*i*.*e*., more similar) to the cell niche embedding. The magnitude of the proximity is controlled by a parameter *p*. A higher value of the parameter *p* gives more weight to nearer subgraph embeddings, while a lower value incorporates a broader range of subgraph embeddings. The supervised learning training objective ensures that subgraph embeddings and, implicitly, cell niche embeddings are arranged to separate positively and negatively labeled samples. This scoring system reveals the spatial tissue patterns prioritized by the model that correlate with each patient group. These spatial tissue patterns range from non-discriminative (CPS between −0.5 and 0.5) to strongly correlated with either phenotypic group (|CPS| *>* 0.5). We distinguish between positively-discriminative CPS scores (‘Positive-CPS’; CPS *>* 0.5) and negatively-discriminative CPS scores (‘Negative-CPS’; CPS *<* −0.5).

#### 2.3.2 Cell-Type Specific Spatial Cell and Marker Patterns

To better understand these spatial tissue patterns, we developed a multi-scale statistical framework to analyze cell niches and identify patterns that distinguish responders from non-responders. For each cell, we quantify its local neighborhood, or niche, by checking whether each cell type is present at each possible hop distance (0-, 1-, 2-, and 3-hops; (**Fig. 3a**). Each cell’s niche is represented by a binary matrix, which we denote as a Typed Cell Niche (TCN), where rows and columns correspond to distances and cell types, respectively. Each matrix entry, which corresponds to a cell type-distance pair (‘CT-Distance pair’), indicates the presence of at least one cell of the corresponding cell type (column) at the specified hop distance (row).

We first group cells whose TCNs share identical binary patterns - that is, cells that have the same presence/absence pattern of cell types at each distance - into what we term “TCN clusters”. To identify TCNs that distinguish phenotypes, here immunotherapy sensitive versus resistant tumors, we analyze the distribution of Positive-CPS and Negative-CPS cell niches in each TCN cluster. We compare this Positive/Negative-CPS distribution within each TCN cluster to the same distribution for cells outside it using Fisher’s Exact Test (**Fig. 3b**). This analysis identifies specific cell niche patterns that significantly associate with treatment response.

For each significant TCN cluster, we identify CT-Distance pairs that significantly contribute to the spatial pattern’s predictive power. This second-level analysis compares the Positive/Negative-CPS distribution of cells corresponding to each CT-Distance pair using Fisher’s Exact Test to compare cells within a significant TCN cluster to cells in all other TCN clusters.

For each significant CT-Distance pair, we identify differentially expressed markers within specific spatial and cellular contexts. This third-level analysis compares the distributions of marker expression between Positive-CPS cells and Negative-CPS cells matching the CT-Distance pair and within the TCN cluster. This multi-level analysis ties together spatial patterns predictive of response to therapy at different biological scales from markers to cells to niches.

#### 2.3.3 Spatial Marker Patterns

To discern key spatial marker patterns indicative of the ground truth phenotype, we employed sCCA to model the relationship between the input data and the cell niche embeddings spaces (**Fig. 4a**). The cell niche embedding space is reflective of the phenotype and has a one-to-one mapping to the cells in the input space. By design, sCCA identifies sparse canonical directions, or weight vectors, that maximize the correlation between the input and cell niche embedding spaces, identifying genes most relevant to the cell niche embedding representations (or indirectly to the phenotype). Our approach involves four distinct sCCA assessments to study marker patterns across a 3-hop neighborhood, delineated by graph distances of 0, 1, 2, and 3. Initially, at graph distance zero, matrix *X*^[0]^ records the transcriptional profiles of individual cells, which undergo sCCA in relation to matrix *C*^[0]^ containing their corresponding cell niche embedding profiles. Progressing to graph distance one, matrix *X*^[1]^ compiles the transcriptome profiles from each cell’s immediate neighbors (orange, **Fig. 4a**). These profiles are then analyzed through sCCA against matrix *C*^[1]^ which duplicates the cell niche embedding profile of the reference cell for each neighbor (pink, **Fig. 4a**), registering each neighbor’s transcriptional profile with the cell niche embedding of its reference cell (blue registration, **Fig. 4a**). At graph distance two, matrix *X*^[2]^gathers profiles from the secondary neighborhood, excluding immediate neighbors, and these profiles are subjected to sCCA against matrix *C*^[2]^ which assigns the cell niche embedding profile of the reference cell to each secondary neighbor. This systematic registration of matrices *X*^[*k*]^ and *C*^[*k*]^ at incremental distances facilitates a multiscale sCCA analysis, providing deeper insights into the spatial and cellular mechanisms at play. Specifically, for each of the four graph distances (0, 1, 2, or 3), we retain five weight vectors, resulting in a total of 20 weight vectors. These vectors are sparse, with their nonzero elements specifying the genes most strongly associated with the cell niche embedding representations. Each marker’s importance is determined by the *ℓ*_2_ norm of its entries across all 20 weight vectors—the larger the norm, the more crucial the marker.

### 2.4 SORBET Identifies Key Spatial Cell Patterns Differentiating Immunotherapy Response from Resistance in Metastatic Melanoma

As described above, to associate the cell niche embedding representation of each cell with the measured phenotype, we compute its Cell Phenotypes Similarity score (CPS; **Fig. 2a**; see 2.3). To visually illustrate the distribution of CPS scores across all cells and across all samples, we employ UMAP [32, 33] and color each cell with its corresponding CPS score (*p* = 3; **Figure 2b**). To identify tissue regions associated with response or non-response to immunotherapy, we overlay the CPS scores for each cell on the sample (**Fig. 2c**).

We highlight four tissues from the metastatic melanoma dataset, two from patients who responded to immunotherapy and two from patients who did not. The CPS score is visualized through a color-bar where red indicates response to immunotherapy, blue signifies non-response, and grey denotes no clear association (**Figure 2a,b**). Typical immunofluourescence techniques, which measure a handful of markers (see **Figure 2c**, bottom), are insufficient to directly identify phenotype-associated tissue patterns. Notably, we observe in some samples a mixture of predicted responsive and non-responsive cell niches in tissue samples derived from non-responding individuals. In more detail, considering CPS scores with absolute value greater than 0.5, allows us to identify regions strongly associated with immunotherapy resistant or sensitive phenotypes. We observe that twelve out of the twenty-seven patients (12/27) not responding to therapy demonstrated a mixture of responsive and non-responsive cell niches. Conversely, only one individual out of twenty (1/20) demonstrated a mixture of predicted responses in the subset of patients responding to therapy. Notably, this patient demonstrated a partial response to therapy, as opposed to a complete response to therapy. Consequently, the predicted mixture of responses may accurately correspond to an intermediate response to therapy phenotype. Similar plots of all 47 tissue samples are included in **Supplementary Figure ??**.

The mixture of positive and negative CPS scores patterns assigned to treatment-resistant tumors aligns with extensive experimental evidence demonstrating a relationship between tumor heterogeneity and poor response to targeted therapy [34–36]. Moreover, biopsies from different tumors in individual patients has demonstrated heterogeneity in the immune milieu and T cell subsets [37]. However, experimental assessment of therapeutic response at the level of cell niches is, to our knowledge, not directly measurable.

To validate that Positive-CPS scores reflect response-associated characteristics even within tissues from patients not responding to immunotherapy, we performed unsupervised, unbiased transcriptional analysis. Again, we split cells into two classes of discriminative CPS scores: Positive-CPS (CPS *>* 0.5) and Negative-CPS (CPS *<* −0.5). We conducted principal components analysis (PCA) on the original marker profiles of cells from either discriminative class, categorizing cells into three groups: Positive-CPS cells from responder tissues, Negative-CPS cells from non-responder tissues, and Positive-CPS cells from non-responder tissues (**Fig. 2d**). The PCA visualization revealed that Positive-CPS cells from non-responder tissues exhibited expression patterns more similar to Positive-CPS cells from responder tissues than to Negative-CPS cells from non-responder tissues. This separation was consistent across the first five principal components (**Fig. ??**).

To quantify this relationship, we examined the k-nearest neighbors (k=50) for each cell in the expression space (**Fig. 2e**). Positive-CPS cells from non-responder tissues showed a high proportion of neighbors derived from responder tissues (mean=45%, median=46%), indicating a distribution spanning both Positive-CPS cells from responder tissues (mean=69%, median=76%) than Negative-CPS cells from non-responder tissues (mean=20%, median=12%). These findings suggest that regions dominated by high CPS scores within non-responder tissues share transcriptional characteristics with responder tissues.

We validated the reliability of the CPS score using *ℓ*1-regularized logistic regression classifiers applied to the problem of identifying cells by response phenotype. For each cell type, we generate two datasets: cells labeled using the original tissue label or cells labeled by CPS score, excluding all cells assigned uncertain CPS scores (|CPS| *<* 0.5). Classifiers were validated using the same approach as our original model; namely, random 5-fold cross-validation with cells split at the patient level, repeated ten times. On average, classifiers using the CPS scores demonstrated superior performance when compared to those using the original label (AUROC = 0.76 using CPS; AUROC = 0.72 using Original Label). This improvement was most pronounced for T cells (AUROC = 0.70 using CPS; AUROC = 0.66 using Original Label) and B cells (AUROC = 0.79 using CPS; AUROC = 0.75 using Original Label). These findings suggest that the CPS score identifies the parts of each tissue most consistent with either label (responder vs non-responder), suggesting that the GNN effectively removes biological signal not related to response to immunotherapy.

#### 2.4.1 Key Cell Patterns Highlight T Cell Activation in Response to Immunotherapy

Next, we analyze the change in CPS scores in the context of well-defined T cell activation processes. T cell activation is partially mediated via a ligand-receptor interaction between the Major Histocompatibility Complex (MHC) and T-cell Receptors (TCR) [38]. CD4-positive (CD4^+^) T cells are activated by professional antigen-presenting cells (APCs) through MHC class II-TCR interaction, while CD8-positive (CD8^+^) T cells recognize antigens presented by MHC class I, which is expressed on most cell types. Here, we use MHC allele expression in cells adjacent to T cells as a proxy for T cell activation. Due to their relevance in cancer immunotherapy[38, 39], we anticipate that activated T cells will be scored differently by our model, when compared with non-activated T cells.

We separately analyzed CD4^+^ and CD8^+^ T cells, and further subtyped CD4^+^ T cells into established subsets (Th1, Th2, Th17, naive) based on their transcription factor expression profiles (see **Sec. 8.7.1**). The CosMx panel included three MHC class I genes (*HLA-A*,*B*,*C)* and seven MHC class II genes (*HLA-DPA1*,*DPB1*,*DQA1*,*DQB1*,*DRA*,*DRB1*,*DRB5*).

For each T cell subset, we examined how their CPS scores related to MHC expression in neighboring cells. For each specific MHC gene, we split T cells into two groups based on whether their neighboring cells expressed or did not express that gene (**Fig. 2f**). Using Mann-Whitney U tests [40], we tested if MHC expression in neighboring cells significantly shifted the CPS score distribution in T cells. P-values were adjusted using the Benjamin-Yekutieli procedure [41].

In the CD4+ T cell subsets, we found distinct activation patterns between Th1 CD4^+^ cells, which primarily demonstrated activation in non-response phenotypes, and naive CD4^+^ T cells, which demonstrated activation with both response and non-response. First, we found that Negative-CPS Th1 cells showed significantly different CPS distributions based on MHC Class II expression in neighboring APCs. Specifically, five of seven MHC Class II alleles in B cells (*HLA-DPB1*,*DQB1*,*DRA*,*DRB1*,*DRB5*, adjusted *p <* 0.05) showed significant associations with Negative-CPS Th1 cells (**Fig. 2g**), consistent with the role of Th1 cells in adaptive immunity. In contrast, expression of a single MHC Class II allele (*HLA-DQPB1*) was significantly associated with Negative-CPS Th1 cells activation (adjusted *p* = 0.010). In contrast, naive CD4^+^ cells demonstrated numerous significant associations with allele expression in adjacent APCs. With B cells, Negative-CPS naive T cells were significantly associated with all MHC Class II alleles and Positive-CPS naive T cells were associated with four of seven alleles (*HLA-DPA1, HLA-DPB1, HLA-DRA, HLA-DRB1*; adjusted *p <* 0.05). In contrast to Th1 cells, naive CD4^+^ T cells demonstrated more and stronger associations with adjacent macrophages including association with expression of three alleles for Negative-CPS cells (*HLA-DQB1, HLA-DRB1, HLA-DRB5*) and three with Positive-CPS cells (*HLA-DRB1, HLA-DRA, HLA-DQA1*). Th2 CD4^+^ T cells were largely not associated with any alleles (one significant association, Negative-CPS Th2 cells with HLA-DRB5 expression in macrophages; **Fig. ??**). These findings suggest a distinct Macrophage-Naive CD4^+^ T cell interaction indicative of response to immunotherapy.

For CD8^+^ T cells, we analyzed CPS distributions in relation to MHC class I expression in neighboring endothelial cells, cancer-associated fibroblasts (CAFs), and malignant cells **Fig. 2h**). We found that CD8^+^ T cells primarily demonstrated associations between the Negative-CPS subset of T cells and MHC Class I expression including two alleles expressed in CAFs (; adjusted *p <* 0.05), one allele expressed in malignant cells (*HLA-B*), and all four MHC class I alleles expressed in endothelial cells. In contrast, only MHC class I alleles expressed in CAFs were associated with response like CD8^+^ T cells (*HLA-A*,*C*,*E*; adjusted *p <* 0.05). CAFs have typically been associated with a tumor-supportive, immunosuppressive role; however, identification of functionally distinct CAF subsets, including tumor-suppressive, immunosupportive antigen-presenting CAFs, have indicated a more complicated role for CAF-mediated adaptive immune system [15, 42].

### 2.5 SORBET Identifies Spatial Cell-Type Patterns Defining Response to Immunotherapy Shared Across Tissues

To identify multi-scale spatial patterns associated with response to immunotherapy, we apply our TCN analysis, as previously described (2.3, **Figure 3a,b**). This analysis identified significant cell niche clusters with shared cell type patterns. With each cluster, we further identify significant cell type and marker patterns. A representative analysis of a non-responder-associated TCN illustrates this multi-scale analysis (**Figure 3c,d,e**), revealing coordinated patterns of CD8+ T cells and B cells at varying distances from malignant cells. All significant TCN clusters, cell types, and marker patterns are included in the supplementary materials (**Supplemental Figures: ??, ??, ??, ??**).

To ensure robust comparisons and statistical power, we filtered cell niches to include those: (1) centered on a ‘Malignant’ cell and (2) are present in tumors from at least 10 patients with at least 50 cell niches in the cluster. For marker analysis, we analyzed only markers that were: (1) expressed in at least half of Positive-CPS or Negative-CPS cells and (2) expressed in at least 1% of total cells, where expressed here is defined as having non-zero counts. These constraints limit the total number of cells analyzed using TCN patterns (2,499 cell analyzed out of 65,615 total cells).

As previously mentioned, the TCN analysis may be computed with either the CPS score or each tissue’s original label. A detailed comparison of the patterns inferred by each methods is beyond the scope of this paper. We quantify the similarity using the Ranked Biased Overlap (RBO) score (**Figure 3f**). RBO measures the similarity between two ranked lists [43] —in this case, the CPS- and label-derived lists of significant markers for each cell type ranked by absolute z-score. A high RBO score indicates strong similarity between the two lists, while a low score suggests greater divergence. Rare cell types in the dataset, such as Endothelial (‘Endo’) and Cancer-associated Fibroblasts (‘CAF’), show the largest differences between scores. These findings suggest that the CPS score may provide stronger insight into rare cell types, when compared with the original tissue label.

Next, we analyze significant marker expression patterns across previously identified TCN clusters associated with treatment outcomes (**Figure 3h**). Here, significant markers are up-regulated consistently in either responders or non-responders across TCN clusters. Consistent with our findings above, responders show high expression of multiple *HLA* alleles —including, *HLA-B, HLA-C, HLA-DRA, HLA-DQA1, HLA-DPA1, HLA-DRB1*, —and *HLA-E*, underscoring robust antigen presentation [44–46]. Additional markers significantly associated with responders implicate enhanced cytotoxic T-cell activity (*IGKC, CD74*) [38, 47, 48] and interferon-*γ*-mediated MHC-I/II expression (*STAT1*) [49].

Conversely, non-responders exhibit elevated *MZT2A, VIM, HSP90AB1, HSP90B1, MALAT1*, and *APOD*, mirroring the metabolic reprogramming, stress responses, and immune suppression characteristic of poorly immunogenic tumors [50–55]. Newly identified ribosomal proteins (*RPL34, RPL37*) are also upregulated in non-responders. While the exact roles of these proteins have not been fully characterized in metastatic melanoma, both *RPL34* and *RPL37* have been implicated in tumor growth and proliferative signaling in other cancers, often via the MDM2–p53 axis or PI3K/Akt pathway [56–58].

Next, we analyzed the changes in significant marker expression patterns of the central malignant cells and surrounding CD8^+^ T cells at increasing hop distances (**Figure 3i-j**). In responders, central malignant cells prominently express *HLA-B, HLA-C, HLA-DPA1, HLA-E, HLA-DRA, HLA-DRB1, IGKC*, and *IF27* indicative of robust antigen presentation and interferon-driven immune activation [38, 44, 59]. Surrounding CD8^+^ T cells show increased *IL32* at hop 2, which is known to enhance dendritic cell and macrophage activation, ultimately promoting T-cell infiltration and synergy with checkpoint blockade [60]. Moreover, responder CD8^+^ T cells upregulate MHC genes, *CCL5*, and *IGKC*, reflecting enhanced T-cell recruitment, robust antigen presentation, and beneficial plasma cell involvement, respectively [61–63]. Specifically, **(author?)** [63] demonstrates how MHC class II–expressing CD8^+^T cells can directly control tumors, **(author?)** [62] shows that *CCL5* promotes effector T-cell migration into melanoma lesions, and **(author?)** [61] highlights an essential role for *IGKC* in plasma cells predicting positive immune-checkpoint blockade responses.These patterns underscore a tumor microenvironment (TME) enriched for cytotoxic activity and immune priming, consistent with improved therapeutic outcomes.

Among non-responders, central malignant cells highly express *MZT2A, MALAT1, GLUL, VIM, HSP90AB1*, and *APOD*. These markers promote proliferation and invasion (*MZT2A* [50]), drive metastatic reactivation and immunosuppression (*MALAT1* [54]), support glutamine-dependent metabolism (*GLUL* [64]), facilitate epithelial-to-mesenchymal transition (*VIM* [65]), and foster stress adaptation and T-cell resistance (*HSP90AB1* [66]), while *APOD* associates with poor outcomes ([55]). Moreover, CD8^+^ T cells at hops 1–3 also upregulate *MZT2A* and *MALAT1*: elevated *MZT2A* correlates with active mitosis and partial exhaustion [67], and *MALAT1* drives short-lived effector differentiation at the expense of durable memory responses [68]. Together, these signatures reveal a dysfunctional microenvironment in non-responders, where tumor stress adaptations and compromised T-cell responses undermine checkpoint blockade efficacy.

These TCN-based findings extend our understanding of spatial organization in immunotherapy response beyond previously reported cell-cell interaction analyses [13, 14, 16]. By integrating cell type patterns with molecular signatures across defined spatial scales, we reveal coordinated tissue organization patterns that may influence treatment outcomes. The consistency of these patterns across patients suggests potential therapeutic strategies targeting specific cellular neighborhoods.

### 2.6 SORBET Identifies Key Spatial Marker Patterns Differentiating Immunotherapy Response in Metastatic Melanoma

To identify spatial marker patterns associated with treatment response, we employed sparse canonical correlation analysis (sCCA) between the input molecular profiles and cell niche embeddings across spatial scales (**Fig. 4a**; 2.3). For each graph distance (0-3 hops), we computed five leading sparse components, creating a marker program matrix (20 components × 960 markers) that captures spatial-molecular relationships.

Ranking markers by their overall contribution, which is calculated as the norm across all components, revealed key determinants of immunotherapy response (**Figure 4b**). The highest-ranked markers included immune-related genes, notably *HLA-A, HLA-B, HLA-C*, and, the gene that encodes the kappa light chain constant domain, *IGKC*, all known predictors of immunotherapy response [38]. Cell division marker *MZT2A* and mesenchymal differentiation marker *VIM* showed the strongest influence. While *MZT2A*’s role in NSCLC metastasis is established [50, 51], its possible association with melanoma progression warrants investigation. Example marker programs across spatial scales illustrate these patterns (**Figure 4c**.

To validate the biological relevance of sCCA-identified markers, we retrained SORBET using only markers with non-zero weights in our analysis (**Figure 4d**. While the performance decreased compared to the full model (AUROC = 0.84 vs 0.95), it significantly exceeded that achieved using excluded markers (AUROC = 0.68), confirming the biological significance of identified markers.

We next sought to understand the spatial basis of the the sCCA-inferred, critical markers. Specifically, we analyze the local spatial correlation of each marker, individually, around each cell using the Local Moran’s I index (8.7.3). The standard (non-Local) Moran’s I quantifies co-variation across an entire graph. In contrast, the local metric quantifies the co-variation between a node, or cell, in the graph and its neighboring nodes for each analyzed marker. Here, we quantify co-variation between each cell and its three-hop neighborhood. This choice mirrors the graph convolution used in the Learning step to compute each cell niche embedding. Within each niche, markers are identified as strongly co-varying —either positively, which we term ‘Clustered’, or negatively, which we term ‘Dispersive’ —or not co-varying. Statistical significance of these associations was estimated using a permutation test (Benjamini-Yekutieli correction [41]). For each marker, this test evaluates computed local Moran indices by comparing between observed and randomly distributed spatial marker expression.

For most markers identified in the sCCA analysis, we identify a large number of cells (*N >* 2000) with substantial local co-variation (**Figure 4e**; ‘Non-responders - Cluster’, ‘Responders - Cluster’). In contrast, markers not identified in the sCCA analysis, on average, include many fewer cells (Responders mean = 207.1, Non-responders mean = 131.1; ‘Non-signif. (Avg.)’). Many markers, including *LGALS3, HSP90AB1*, and *HSP90AA1*, demonstrate more than 1,000 more cells identified in responders, when compared with non-responders. Although these genes are associated with aggressive cancers, both *HSP90* and *Galectin-3* have been proposed as targets to enhance immunotherapy response [66, 69]. These findings show that markers exhibiting strong local co-variation across many cell niches can be identified by the GNN model and distinguish responders from non-responders.

### 2.7 Extending SORBET to Spatial Proteomics of Diverse Pathological Contexts

To test SORBET’s applicability across different diseases and orthogonal high dimensional spatial molecular platforms, SORBET was applied to two previously reported spatial proteomics datasets. These datasets included samples from Non-Small Cell Lung Cancer (NSCLC) [70] and Colorectal Cancer (CRC) [19] (**Table ??**; 8.8.2, 8.8.3). The first NSCLC dataset included 29 primary tumors represented in tissue microarrays. Protein abundances for 29 markers were quantified using Imaging Mass Cytometry (IMC). Patients received PD-1 / PD-L1 blockade therapy and their response to treatment was recorded (11 responders, 18 non-responders). The classification task for the NSCLC dataset is to distinguish responders from non-responders. The second CRC dataset consisted of 35 patient-derived tissues with 49 protein markers profiled using the CO-Detection by indEXing (CODEX) technique. The CRC samples were classified based on histologic structures into Diffuse Inflammatory Infiltrate (DII) and Crohn’s Like Reaction (CLR), the latter being indicative of favorable prognosis. The classification task for the CRC dataset is to distinguish between these two histologic structures.

SORBET significantly outperformed all previously reported baseline methods on the task of classifying NSCLC response to immunotherapy, achieving an AUROC of 0.96 (**Fig 5a**). This performance surpassed that of spatial methods like CytoCommunity (AUROC = 0.71) and SPACE-GM (AUROC = 0.78), as well as non-spatial methods such as CITRUS-SC (AUCROC = 0.75), and Augur-SC (AUROC=0.77). Similarly, SORBET excelled on the CRC tissue structure classification task, attaining an AUROC of 0.98 (**Fig 5b**). This performance surpassed that of the non-spatial methods (CITRUS-SC:0.86, Augur-SC:0.91). Notably, SORBET also outperforms both spatial methods (CytoCommunity: 0.78, SPACE-GM: 0.81). The performance of the latter two methods are concordant with their previously reported results on the CRC dataset. These results showcase SORBET’s robustness across various classification tasks and enhanced predictive ability relative to methods that do not utilize the full omics or spatial information.

**Fig. 5.**
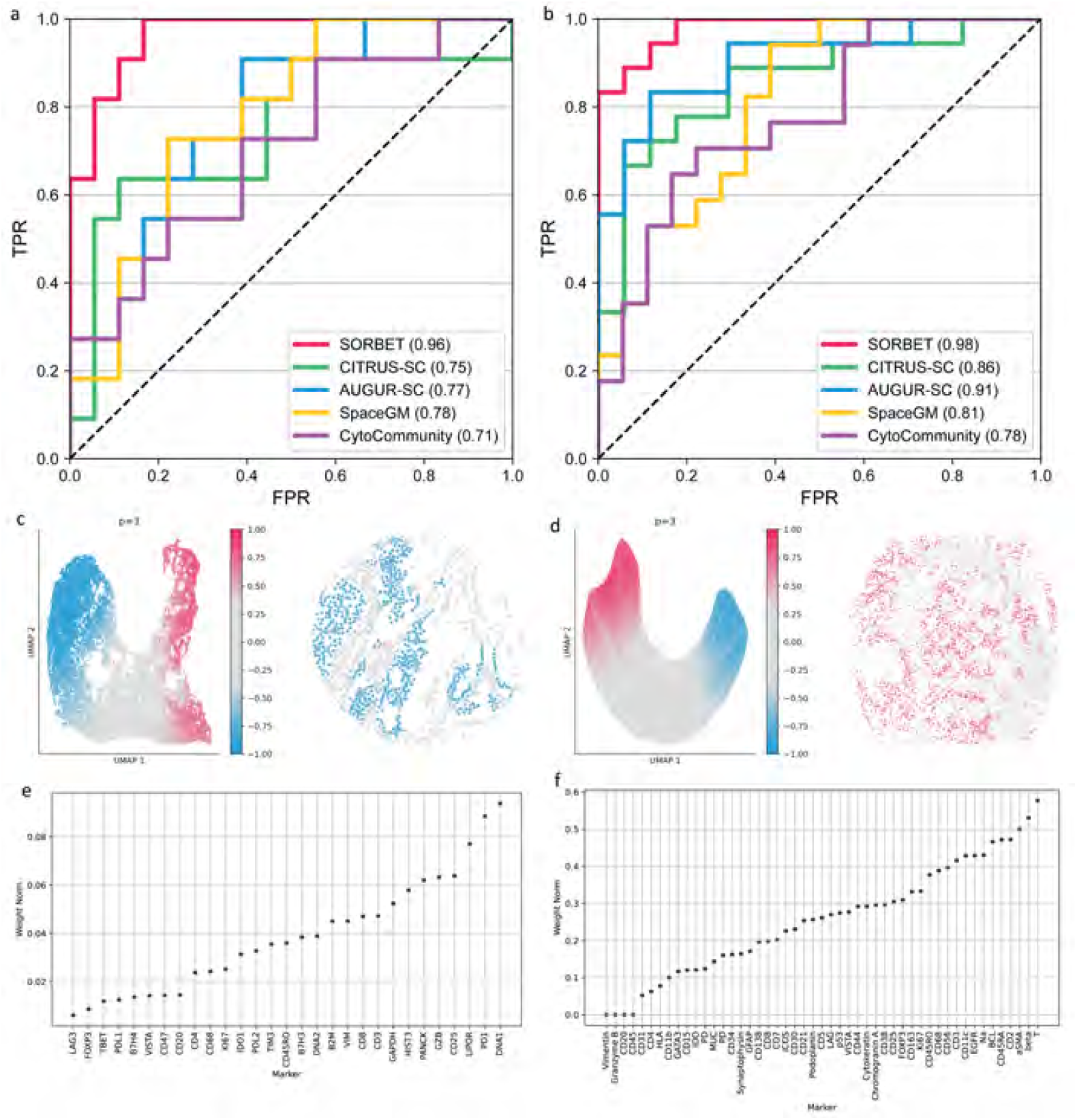
Extending SORBET to Spatial Proteomics Datasets. **(a,b)** SORBET’s performance on the **(a)** NSCLC (IMC) and **(b)** CRC (CODEX) datasets, compared against spatial (SpaceGM, CytoCommunity) and non-spatial (AUGUR, CITRUS) methods, using 5-fold cross-validation with ten random re-initializations, as detailed in **Figure 1a. (c,d)** Key spatial cell patterns for **(c)** NSCLC and **(d)** CRC datasets, as shown in **Figure 2a-b**. (left) UMAP of the CNEs from each dataset colored according to their corresponding CPS scores. (right) Representative tissues of **(c)** non-response to immunotherapy and **(d)** the CLR histologic type. **(e,f)** Marker importance for the **(e)** NSCLC and **(f)** CRC datasets, corresponding to **Fig. 4a**.

The interpretability approaches employed by SORBET are not confined to spatial transcriptomics datasets. Our methods are directly applicable to spatial proteomics tasks, facilitating insights across spatial technologies. The Cell Phenotypes Similarity score (CPS)-based analysis of tissue samples, as in **Fig. 2**, reveal small regions in tissue samples most correlated with each measured phenotype in both the NSCLC (**Fig 5c**) and CRC (**Fig 5d**) datasets. A complete analysis is available for both the NSCLC (**Supplementary Figure ??**) and CRC (**Supplementary Figure ??**) datasets.

As in **Section 2.6**, we applied SORBET’s approach to identify key spatial marker patterns to analyze the NSCLC and CRC datasets. In the NSCLC dataset, markers including *PD1* and DNA1 (a marker of DNA content) were associated with response to treatment **Fig 5e**)). The analysis also identified several immune markers, *CD25, CD3*, and *CD8*, as key predictors of response to PD1/PDL1 therapy. The identification of known tumor biomarkers (*i*.*e*., *PD1* [71–73]) and immune markers [74–76] identified here is consistent with known biology underlying immunotherapy-treated tumors. A re-analysis of the CRC dataset (**Fig 5f**) identified markers associated with lymphocyte differentiation, including *CD2, CD45Rα*, and *CD45RO* [38], as important discriminators of the DII and CLR phenotypes. These findings are consistent with previous studies, which have demonstrated the importance of lymphocyte differentiation for evaluating CLR lymphoid structures in CRC [77, 78].

## 3 Discussion

SORBET, a geometric deep learning framework, integrates spatial information and complete omics data to facilitate effective classification of multiple samples by observed phenotype. Moreover, interpretability methods developed specifically for biological phenotype prediction reveal the spatial molecular and cellular patterns underlying this classification. We demonstrate our findings across several spatial cellular profiling technologies, including CosMx, IMC and CODEX, and across different disease contexts and phenotypes. The combined methods for both precise classification and detailed biological insight underlines the framework’s potential to drive forward personalized medicine and biomolecular research.

We demonstrated that SORBET facilitates effective classification of noisy, high-dimensional data with limited sample sizes. To improve performance, we applied a novel automatic subgraph extraction algorithm to the data. This data augmentation strategy permits deep learning analyses of small datasets (n = 29 (NSCLC), 35 (CRC), 47 (Melanoma)). SORBET exploits whole cell profiles, capturing the information of diverse cell states and their spatial contexts in relation to observed phenotypes. We showed that our proposed method is superior to both non-spatial, single-cell classification methods (CITRUS, Augur) and published spatial proteomics GNN-based deep learning methods (SPACE-GM, CytoCommunity). Previous geometric deep learning methods, such as SPACE-GM and CytoCommunity, limit the input data complexity using cell-typing, which may discard important functional and biological information. Critically, our work is, to our knowledge, the first to effectively analyze spatial transcriptomics data.

The unique architecture of SORBET is tailored to unveil phenotype-dependent structures and patterns across different biological states and scales —from markers in the sCCA method to cell niches using Cell Phenotypes Similarity scores with numerous detailed patterns captured in the TCN analysis. While the GNN captures complex, non-linear spatial marker patterns, the interpretability methods used here map these representations into linear patterns, highlighting a persistent challenge in the field of deep learning interpretability. Nevertheless, our findings are consistent with established tumor immunobiology, providing strong evidence that the GNN uncovers meaningful biological patterns. We anticipate that our interpretability approaches will provide rich new lines of inquiry into the diverse, complex mechanisms underlying response to immunotherapy. Moreover, the described methods are novel technically and applicable to most GNN-based models.

At present, available spatial omics datasets have limited numbers of samples. Consequently, models developed for these datasets are typically of limited complexity and have limited potential for generalization. We anticipate that emerging spatial omics data will facilitate extension of our methods outside of cancer-associated binary classification contexts to other biological contexts and to multi-class or continuous outcomes, such as Progression-Free Survival. Furthermore, independent validation on matched datasets would provide additional confirmation of model efficacy.

In summary, we demonstrate the potential SORBET has to offer for a broad range of applications. For biologists, SORBET’s interpretability methods enable identification of spatial molecular and cellular patterns suitable for further validation and targeting. For clinicians, especially pathologists, SORBET demonstrates superior classification of patients into different treatment groups. Currently, correlating spatial omics with observed phenotypes is hard. SORBET has the potential to bridge this gap, delivering interpretative insights that seamlessly connect spatial molecular and cellular patterns to the biological processes they orchestrate.

## 4 Data and Code availability

The NSCLC and CRC datasets were previously published. The Metastatic Melanoma dataset will be published pending paper publication. The source code for SORBET will be made available on paper publication.

## 5 Author Contributions

SS, JMC, RT and YK conceived the study. YK provided overall supervision of the study. RT, KS, and HK provided additional supervision. KS provided the NSCLC samples with SJD generating the resulting IMC dataset. DR and HK provided metastatic melanoma samples and clinical data. LA curated clinical annotations and the metastatic melanoma TMA. HK and YK provided the CosMx melanoma dataset. SS and JMC developed the model architecture and interpretation framework, wrote the final code and performed the data analysis. SS, JMC, and YK wrote the manuscript with input from all co-authors.

## 6 Acknowledgments

YK discloses support for the research of this work from NIH [R01GM131642, UM1PA051410, U54AG076043, U54AG079759, U01DA053628, and R33DA047037], HK discloses support for the research of this work from P50CA121974. KAS discloses support for the research of this work from R37CA245154 and R01CA262377.

## 8 Methods

### 8.1 Input Data

SORBET is a framework that classifies tissue samples by observed phenotypes, represented as binary labels. Tissue samples may be profiled using multiple spatial omics platforms, including spatial transcriptomics and spatial proteomics approaches. The framework assumes that the data comprise cells with spatial annotation and assigned multiplexed features. Preprocessing steps to identify cell boundaries, deconvolve overlapping cells, quantify measured markers and associated technical challenges (*e*.*g*. [79–83]) are not the explicit focus of this work.

Input data are tissues, 𝒯^(*i*)^ = (𝒳^(*i*)^, *𝒞*^(*i*)^ |ℰ ^(*i*)^), and binarized labels, *y*^(*i*)^ ∈ {0, 1}. First, each tissue is defined by cells profiling data, 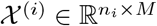, where *n*_*i*_ denotes the number of cells in the tissue, 𝒯^(*i*)^, and *M* denotes the number of profiled markers. Second, each tissue is defined by either cell spatial locations, 𝒞^(*i*)^ ∈ ℝ^*n×*2^, or by predefined edges, ℰ ^(*i*)^ = *{*(*j, k*) | *j, k* ∈ *{*1, 2, …, *n}*; *i* ≠ *j}*. A dataset comprising *N* tissues is denoted 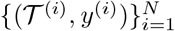. We call the task of predicting the label, *y*^(*i*)^, from the tissue, *T* ^(*i*)^, tissue classification.

### 8.2 Graph Modeling

To translate tissues, 𝒯 ^(*i*)^, to graphs, 𝒢^(*i*)^, we must identify appropriate vertices and edges. Each cell is treated as a vertex, 𝒱^(*i*)^ = *{*1, 2, …, *n}*, with arbitrary indexing. Datasets may have predefined edges ℰ^(*i*)^ where the edge (*j, k*) ∈ ℰ^(*i*)^ if the cells *v*_*j*_, *v*_*k*_ ∈ *𝒱*^(*i*)^ are identified as having a physical contact. Otherwise, edges may be inferred from input cell locations 𝒞^(*i*)^, where an edge (*j, k*) ∈ ℰ^(*i*)^ if the Euclidean distance 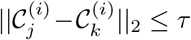 for some tunable distance threshold *τ*.

With vertex indexing and an edge set, either predetermined or inferred, a graph is defined as 𝒢^(*i*)^ = (𝒱^(*i*)^,ℰ^(*i*)^, 𝒳^(*i*)^). The graph construction is independent of downstream analysis. Numerous biologically reasonable graph construction methods might replace this choice.

### 8.3 Cell Typing

In our study, cell typing was conducted by integrating spatial transcriptomics data from NanoString’s CosMx platform with the annotated single-cell RNA sequencing reference dataset GSE115978 [84]. This dataset, derived from 31 melanoma tumors, focuses on the cellular landscape relevant to immunotherapy resistance and includes detailed annotations of various cell types, crucial for understanding clinical response dynamics. It is publicly available and can be accessed via NCBI GEO.

Raw counts, metadata, and field-of-view (FOV) information were processed using the Squidpy[85] and Scanpy [86] libraries. During preprocessing, cells with fewer than 100 total gene counts and genes detected in fewer than 400 cells were filtered out to ensure high-quality data. The remaining data underwent normalization and log transformation to stabilize variance and improve comparability.

For cell type classification, we employed the scVI model [87], training it on concatenated spatial and reference datasets. The scVI model was set up on a GPU for efficient processing, with a batch size of 128 and a training duration of 100 epochs to balance computational efficiency and model performance.

Cell type predictions were made using a nearest-neighbor classification approach in the latent space derived from scVI. We used a nearest neighbors algorithm with *k* = 1 to ensure each spatially resolved cell was mapped to the single closest reference cell type based on their latent representations. This precise approach, along with detailed hyperparameters, data handling steps, and the utilization of a well-curated reference dataset, has been provided to ensure reproducibility and fidelity in cell typing across similar studies.

### 8.4 Automatic neighborhood extraction

Next, input graphs, 𝒢^(*i*)^, were automatically processed to extract relevant subgraphs, 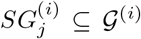. SORBET is applied to subgraphs for two reasons. Biologically, subgraphs provide multiple, localized views of tissues that capture heterogeneity within tissues. Technically, this choice augments available data improving performance and robustness of classifiers. In this work, the subgraph extraction algorithm increases the dataset size by an order of magnitude.

A subgraph 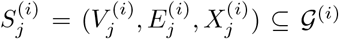, is defined on a subset of the vertices, 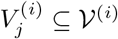. Edges and marker data are filtered to include only vertices 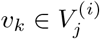, such that 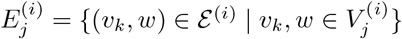. Consider a function, *f*: *v*_*j*_ ↦ ℝ for *v*_*j*_ ∈ 𝒱 ^(*i*)^, that maps a cell to some real value. The algorithm (**Alg. 1**) identifies ‘anchor cells’, or cells prioritized by the function *f*, and extracts a subgraph around each anchor cell. For each anchor cell, the median of *f* is taken on a one hop neighborhood to ensure the cell is not an outlier. The anchor cell is used if the median on the 1-hop neighborhood exceeds the median across all cells. Subsequently, a subgraph is formed by taking the *k*-hop neighborhood around the anchor cell if the neighborhood is of minimum size *MS*. For this work, we choose *k* = 10 and *MS* = 50. This procedure identifies a set of subgraphs 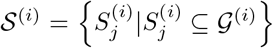 where 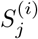 denotes the identified subgraph anchored at vertex, *v*_*j*_ ∈ 𝒱^(*i*)^. We associated the graph label, *y*^(*i*)^, with each extracted subgraph. Consequently, the sub-tissue classification task comprises the dataset 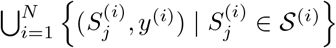.

#### Algorithm 1 (Automatic Subgraph Extraction)

The function med denotes the median. 𝒩_*k*_(*v*) denotes the *k*-hop neighborhood around a chosen cell, *v*. Define 𝒢(*i*) and *f* as above.

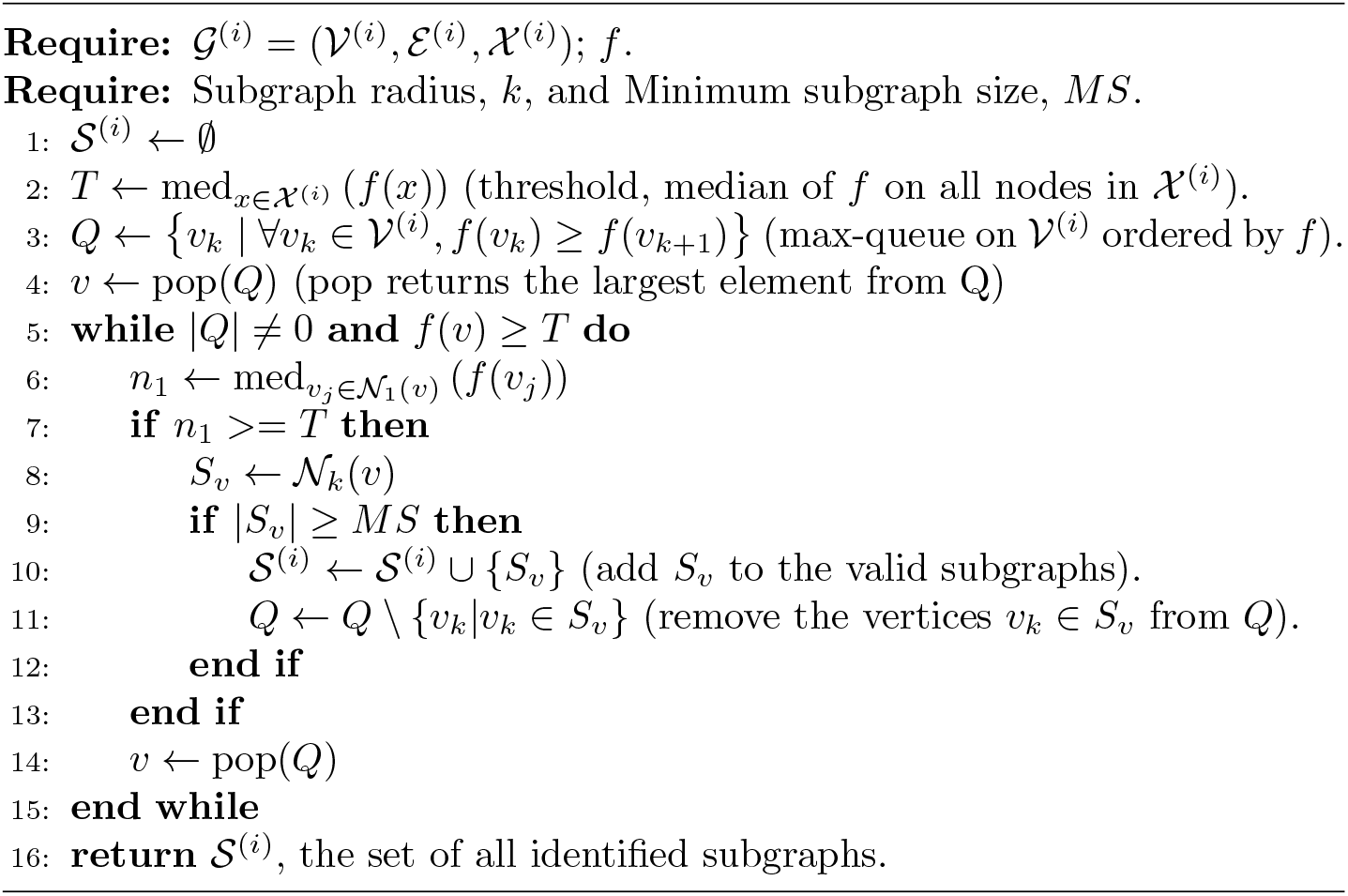

The algorithm’s output depends on the chosen function, *f*, and the graph connectivity. Here, we choose a biologically relevant marker (*e*.*g*., an epithelial cell marker such as *S100B*) and define *f*: *v*_*j*_ ↦ *x*_*j*_(*m*) where *x*_*j*_(*m*) denotes the value of the marker *m* measured for cell *v*_*j*_. Note, *f* is not constrained to adapt a marker value; it may take any relevant value (*e*.*g*., node degree, centrality, *etc*.). Moreover, a random selection of subgraphs may be chosen if no prior information is available.

### 8.5 SORBET Model Architecture

We propose a graph convolutional network (GCN) based neural network architecture to analyze the graph modeled tissues and their corresponding sub-graphs. For discussion, the GCN network may be broken into four modules: an embedding module, a GCN module, a cell niche embedding module, and a subgraph prediction module. We use the superscripts (*e*), (*g*), (*c*), and (*s*) to denote variables pertaining to these four modules, respectively. Each module may comprise multiple layers; here, we use *ℓ*_0_ to denote the input to the module, and *ℓ*_max_ to denote the output. As discussed above, each tissue sample in the tissue classification task is broken into subgraphs, 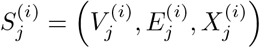, where *X*^(*i*)^ ∈ ℝ^*n×M*^ is the subset of omics profiles for the 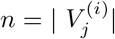 vertices present in the subgraph.

The model architecture predicts a label for each subgraph as follows. First, the embedding module transforms the *M*-dimensions at each channel into a rich intermediate representation using a multi layer fully-connected neural network (FCNN). The FCNN structure uses layer normalization (LN) and the Rectified Linear Unit (ReLU) to compute each intermediate representation. At each layer of the FCNN, *ℓ*, a matrix 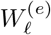 of weights is learned that maps the previous representation, 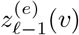, of vertex *v* to the representation 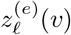. The initial representation, 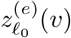, is simply the profiled cell data, 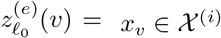. For brevity, we drop the typical superscript, (*i*), denoting the tissue of origin. Formally, the *ℓ*-th layer may be written

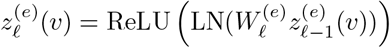

where LN denotes the layer norm and ReLU denotes the ReLU non-linearity. The layer norm, LN, is defined as 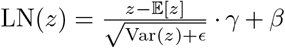 where 𝔼 and Var denote the expectation and variance, respectively. The parameters *γ* and *β* are learned parameters and *ϵ* = 1*e* − 5 (default) [88]. The ReLU non-linearity is defined as ReLU(*z*) = max(0, *z*). We denote the final representation obtained by the embedding module as 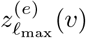.

Next, the GCN module combines Deep GCN layers connected with dense feed-forward connections, as suggested in [23, 24] and implemented in [89]. Briefly, this module infers weights, 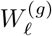, that compute the messages passed among adjacent vertices. Again, we denote intermediate representations of vertex *v* at layer 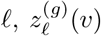. Note, the input representation, 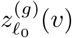, is equal to the output of the embedding module, 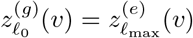. Each layer in the GCN module computes the function,

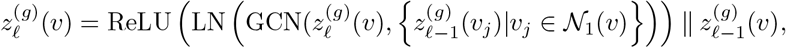

where GCN denotes the standard graph convolutional layer, ∥ denotes concatenation of two vectors (*i*.*e*. a residual block connection), and LN and ReLU as previously defined. As above, we write the final representation 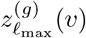

Next, the cell niche embedding module takes the inferred representations of each node and further compresses the data. This module follows the same structure as the embedding module. Specifically, it is an FCNN with the same model architecture (layer normalization and the ReLU activation function). Note, the choice of hyperparameters (*e*.*g*., number of layers and layer width) are unique to each module. Explicitly, the input is 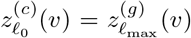, weight matrices, 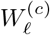, and intermediate representation 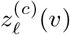, which may be written,

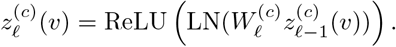

As above, the final representation is noted by 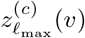 Note, 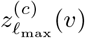 is the cell niche embedding (CNE) of the cell niche centered at *v*.

Finally, the subgraph prediction module applies a global pooling operation, `max-pool`, and a final FCNN. Specifically, the subgraph prediction modules pools over all vertices 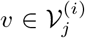 along each dimension. Note, input representation, 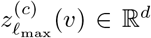, is a vector representation of a cell’s CNE where *d* is a network hyperparameter. We denote the *k*-th vector index of a vector *z, z*(*k*). The `max-pool` operation combines the vector representations, 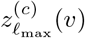, of each 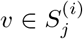 into a single subgraph representation, 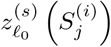, for each subgraph, 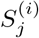. Explicitly, the initial subgraph representation is written

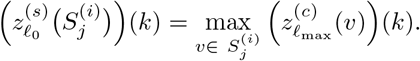

Note, 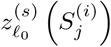 is the subgraph embedding (SE) of the subgraph 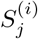.

The SE is passed through a final FCNN with the same structure as the embedding and cell niche embedding modules (layer normalization, ReLU). As above, the intermediate representations may be written

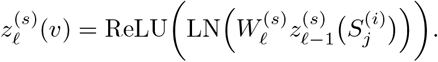

The final layer replaces the ReLU non-linearity with a sigmoid function, *σ*(*z*) = (1 + *e*^−*z*^)^−1^,

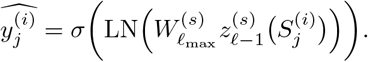

By construction, this prediction is the model’s confidence that *y*^(*i*)^ is 1 (*i*.*e*., positively labeled) given the subgraph, 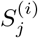,

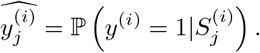

Here, 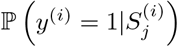 denotes conditional probability in the standard way. The final tissue prediction, ℙ(*y*^(*i*)^ = 1|𝔾 ^(*i*)^), is computed as the mean of the predictions over all of the tissue’s subgraphs,

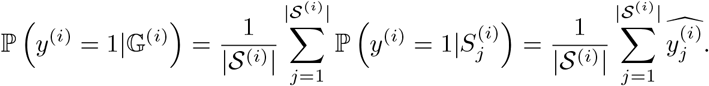

### 8.6 Learning Step - SORBET

In this study, we used a stratified 5-fold cross-validation approach to evaluate the performance of our model. Each dataset was split into five groups (approx. 20% of the data) with positively and negatively labeled samples equally split across each fold. For each held-out test fold, appropriate model structure was identified on the remaining folds – one fold for validation and the remaining folds for training. Model structure was optimized using the Ray tune parameter optimizer. Model trials were limited to 100 hyperparameter combinations. Hyperparameters were set to a single set of values across experiments: 50 epochs, 3e−4 learning rate, and 8 batch size. Models were optimized using the Adam optimizer (default aside from learning rate).

This structure differs from traditional approaches to *k*-fold cross-validation. Typically, three folds might be held-out – two folds to fit model structure and hyperparameters, respectively, and one fold to test. We chose to optimize only model structure (*i*.*e*. layer size, depth) due to limited available data. Additional hyperparameter tuning would likely improve test set accuracy.

Due to the small sample size, we repeated 5-fold cross validation ten times on ten random data splits (50 total models, five-fold cross validation repeated ten times). Model performance is reported using the receiver operator curve (ROC). Two metrics are returned, the mean area under the ROC curve (mAUROC) (*±*standard deviation) and the mean ROC curve (mROC) (*±*standard deviation). Let *g*(*y, ŷ, z*) denote the estimated true-positive rate according to the ROC curve for labels *y* and predictions *ŷ* at a false-positive rate *z*. The mAUROC is computed over the ten random splits, *j* = 1, …, 10,

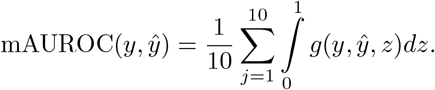

Note, the inner integral is simply the typical AUROC; the mean is taken over the ten random splits. The mROC is computed point-wise for 1,000 points uniformly split in the interval [0, 1]. At each point, let

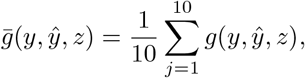

where 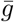 denotes the mROC curve.

For comparison, we consider four methods: CITRUS [26], Augur [27], SpaceGM [14] and CytoCommunity [16]. CITRUS and Augur are methods developed for analysis of spatial transcriptomics without consideration of spatial context. Throughout the text, we append ‘-SC’ to CITRUS and Augur to differentiate these methods developed from methods that consider spatial context. SpaceGM and CytoCommunity are GNN-based methods developed to analyze spatial proteomics datasets. Both methods were previously applied to the CODEX CRC dataset. We have adapated each methods to analyze the metastatic melanoma and NSCLC datasets. For evaluation, we employed the default settings and hyperparameters provided by the methods.

### 8.7 Reasoning Step - SORBET

SORBET provides a natural mechanism for understanding the predictions that were inferred during the Learning Step. In particular, SORBET’s Reasoning Step captures two aspects of the problem: phenotype-dependent tissue architecture and phenotype-dependent cell marker profiles. This step analyzes three embedding spaces, the spatial omics input data space, the inferred cell niche embedding (CNE) space, and the subgraph embedding (SE) space. We note that the analyses methods shown here are are general and novel deep learning explainability techniques. We believe that these may be of general interest to the machine learning community.

#### 8.7.1 Spatial Cell Pattern Analysis

As described in Section 8.6, SORBET employs a `max-pool` pooling operator to calculate a single embedding for each subgraph from the cell niche representations. Subsequently, it predicts the likelihood of that subgraph embedding, which we term the SE, for the observed phenotype. -¿ The max-pooling quantifies each SE from its own CNEs; you still did not explain how the SEs are combined for the final phenotype prediction; this sentence is vague and confusing. Specifically, this pooling operator maps the CNEs that comprise a subgraph, 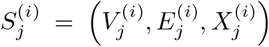, to a single SE. We write each CNE 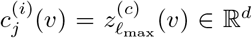, where 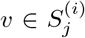 denotes the cell associated with each CNE, 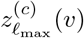 corresponds to the notation in Section 8.6 and *d* is the output embedding dimension of the cell niche embedding. As previously noted, the structure of the pooling operator ensures that the SEs and CNEs occupy the same space (*i*.*e*. demonstrate the same support).

We compute this similarity using an inverse-distance weighting scheme relative to these summary vectors. We term this score Cell Phenotypes Similarity. Briefly, the inverse-distance weighted score function, *w*: *c*(*v*) → [−1, 1], is defined

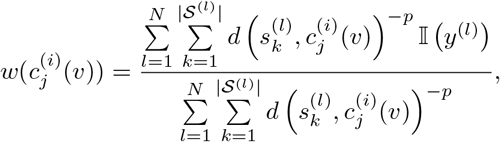

where 𝒮^(*l*)^ is the set of all subgraphs extracted from the graph 𝒢^(*l*)^ (as above), *d* (·,·) is a distance metric, *p* is a chosen scaling parameter, and 𝕀 is an indicator function that maps the binary labels to −1 and 1 (*e*.*g*. 𝕀(0) = −1, 𝕀(1) = 1). Note, *w* ∈ [−1, 1]

Here, *d*(·,·) is Euclidean distance. The choice of value for parameter *p* is reported in the main text. The distance, *d*(·,·), is computed on the leading principal components of the matrix of the CNEs, *C*, where *C* has dimensions *n*^(*c*)^ *×d* where 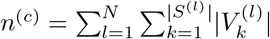 (*i*.*e*., the total number of CNEs across all subgraphs) and *d* is the dimensionality of CNEs. Principal components associated with less than 0.01 explained variance ratio are excluded.

As described, we assess Cell Phenotypes Similarity (CPS) scores by analyzing changes in antigen-presenting cells (APC) adjacent to T cells. First, identify T cells (CD4+ and CD8+) adjacent to appropriate APCs (B cells and macrophages near D4+ T cells; malignant cells near CD8^+^ T cells). CD4^+^ T cells were further typed by well-described sub-types (Th1, Th2, Th17, Tfh, Naive). Cell assignment was based on (binarized) expression of known transcription factors associated with each cell type. CD4^+^ T cells not expressing any transcription factor associated with mature cells were classified as naive. T cells expressing conflicting transcription factors were excluded. For further analysis, we focused only on Th1 and Naive T cells

For each T cell-APC combination, data were first split by response status (responder vs non-responder). APCs were further split by (binarized) expression of a known MHC gene. The studied genes correspond to established MHC-TCR interactions (MHC class II with CD4+; MHC class I with CD8+). A one-sided Mann-Whitney U test was computed between the APCs expressing or not expressing a chosen gene. For example, the MHC class II gene *HLA-DPA1* was studied in the context of CD4^+^ T cells. Its expression was studied in resident Macrophages and B cells both in responders and in non-responders (four total tests). As stated, the chosen CosMx panel captures 10 MHC genes: three class I genes (*HLA-A*,*B*,*C*) and seven class II genes (*HLA-DPA1*,*DPB1*,*DQA1*,*DQB1*,*DRA*,*DRB1*,*DRB5*). In total, we compute 62 statistical tests, six for malignant-CD8^+^ T cell interactions (three MHC class I genes, responders / non-responders), fourteen for macrophage-Th1 CD4^+^ T cell (seven MHC class II genes, responders / non-responders), fourteen for macrophage-naive CD4^+^ T cell, fourteen for B cell-Th1 CD4^+^ T cell, and fourteen for B cell-naive CD4^+^ T cell.

#### 8.7.2 Typed Cell Niche (TCN)

Spatial cell organization was analyzed using Typed Cell Niches (TCNs), where each cell *v* is represented by a binary matrix *T* (*v*) ∈ *{*0, 1*}*^4*×c*^, with *c* being the number of distinct cell types. Each matrix element *T* (*v*)_*i*,*j*_ encodes the presence of cell type *τ*_*j*_ at hop distance *k* = *i*− 1, where *i* ranges from 1 to 4. Define the *k*-shell, *R*_*k*_(*v*), of a cell, *v*, as the set difference between the *k*-hop and (*k* − 1)-hop neighborhoods *R*_*k*_(*v*) = 𝒩_*k*_(*v*) \ 𝒩_*k*−1_(*v*). We set *T* (*v*)_*i*,*j*_ = 1 if there exists 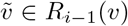 with 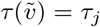, and 0 otherwise, where *τ* (*v*) ∈ [1, …, *c*] defines a map of vertices to the index of their corresponding cell type.

We defined TCN clusters as sets of cells sharing identical neighborhood binary cell type patterns, 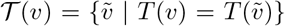, where equality requires 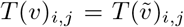 for all indices. Statistical analysis proceeded through three hierarchical levels, with each subsequent level analyzing only patterns found significant in the previous level. At each level, analyses were performed using either CPS scores *w*(*v*) ∈ [−1, 1] or each cell’s source tissue labels *ℓ*(*v*) ∈ {0, 1}.

Pattern-level analysis used Fisher’s Exact tests with contingency tables:

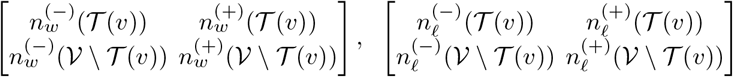

for CPS analysis (left), where 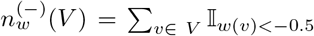 and 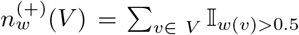, and for tissue label analysis (right) where 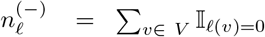 and 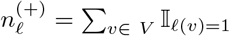. Here 𝒱 denotes the set of all cells in the data, and 𝒱 \ 𝒯 (*v*) is the set of all cells that are not in 𝒯 (*v*).

For each significant TCN pattern, 𝒯_∫_ (*v*), we analyzed cell type-distance pairs, (*τ, k*). Define 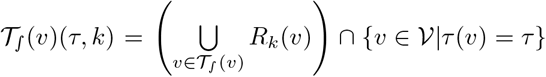 as the set of all cells at hop *k* in cell niches that are part of the TCN cluster 𝒯_∫_ (*v*) (the left term in the set intersection) of cell type *τ* (the right term). Fisher’s Exact tests used contingency tables:

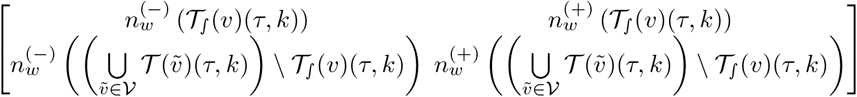

with corresponding tables for tissue labels. The bottom row corresponds to all cells in any TCN pattern matching the same cell type-hop distance pair 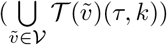, excluding the cells in 𝒯_∫_ (*v*)(*τ, k*).

For significant CT-Distance pairs in each significant TCN, 𝒯_∫_ (*v*)(*τ, k*), we analyzed differences in expression levels for each marker, comparing distributions between positive cells (*w*(*v*) *>* 0.5 or *ℓ*(*v*) = 1) and negative cells (*w*(*v*) *<* −0.5 or *ℓ*(*v*) = 0) within 𝒯_∫_ (*v*) using Mann-Whitney U tests. Each marker *m* may be summarized by a *p*-value, *p*(*m*), which comes from the Mann-Whitney U test, and directionality *d*(*m*) ∈ {−1, 1}, based on the difference in median expression (1 if higher median expression in responders).

For each marker, we report combined statistics across different sets of significant CT-Distance pairs at progressively finer scales:

1. Set constructed across all significant TCN clusters across; across all cell types and all distances.

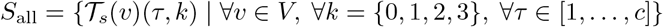
2. For a specified cell type *τ*_*j*_. Set constructed across all significant TCN clusters and all distances.

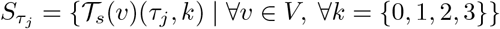
3. For a specified cell type *τ* _*j*_ and distance *k* ^*′*^. Set constructed across all significant TCN clusters

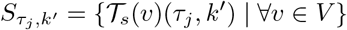

Let *S* denote a generic set from the above types; let *S*[*i*] = 𝒯_*s*_(*v*)(*τ, k*) denote the *i*-th element of *S* (arbitrary indexing). For the significant CT-Distance pair *S*[*i*], let *p*_*i*_(*m*) and *d*_*i*_(*m*) denote the *p*-value and directionality of marker *m*. Using Stouffer’s method, we estimate a combined *z*-score, 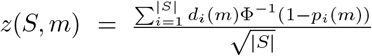, where Φ^−1^ is the inverse cumulative distribution function (CDF) of the standard normal distribution. From *z*(*S, m*), we estimate a combined *p*-value, *p*_*S*_(*m*) = 1 − Φ(*z*(*S, m*)), and combined directionality, 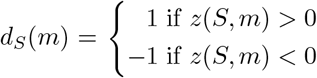. All p-values were adjusted for multiple testing using the Benjamini-Hochberg method, with a false discovery rate (FDR) threshold of 0.05.

#### 8.7.3 Spatial Marker Pattern Analysis

The relative importance of each marker to inform the model’s predictions is assessed by quantifying the shared information between the input data and CNE spaces. By construction, the GCN-based model architecture passes information from all nodes within three hops of a node (*i*.*e*., the cell’s 3-hop niche) back to the node itself. To assess the importance of each marker, we consider the information shared between each node’s CNE and the nodes that pass it information. Here, we adapt a sparse-variant of canonical correlation analysis (sCCA), *ℓ*0-based Sparse CCA, that decomposes the sparse covariance structure between two embedding spaces [25]. This technique allows us to identify the most relevant markers that contribute significantly to the model’s intermediate representations (CNE and SE).

CCA methods are designed to identify linear relationships that maximize the correlation between two paired, or registered, data spaces. The linear relationships form a low dimensional representation of the data spaces. First, we describe the technique for pairing data in the input data and CNE spaces. Let *v* denote a cell under consideration. The CNE associated with *v, c*(*v*) is constructed by convolving the information coming from neighboring cells up to three hops away, *X*_*v*_ = *{x*(*v*_*i*_) | *v*_*i*_ ∈*𝒩*_3_(*v*)}, as well as the node’s intrinsic information, *x*(*v*).

For discussion purposes, we split 𝒩_3_(*v*) into shells, *R*_*k*_(*v*), for *k* ∈ [0, 1, 2, 3]. Here,

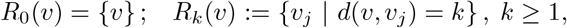

where *d*(*v, v*_*j*_) denotes graph distance. For each shell, *R*_*k*_(*v*), the *d*(*v, v*_*j*_) = *k* exactly. Equivalently stated, each shell is the difference between the *k*-hop and (*k* − 1)-hop neighborhoods, *R*_*k*_(*v*) = *𝒩* _*k*_(*v*) *\ 𝒩* _*k*−1_(*v*) for *k* ≥ 1. The 0-hop shell of a node, the shell with a graph distance of 0, is defined as the node itself.

We construct four sCCA models, one for each hop (*k* = 1, 2, 3) and one for the node’s intrinsic information (*k* = 0). For the intrinsic information, sCCA is computed on the input data space and the output CNE’s where an immediate pairing exists between each central cell’s input data *x*(*v*), and its corresponding embedding *c*(*v*). This pairing procedure involves two matrices, the input data matrix, *X*^[0]^, and the CNE matrix, *C*^[0]^. Each row in the input data matrix, 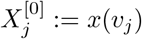, is the omics profile associated with one cell. The corresponding row in the CNE matrix, 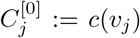, is the CNE associated with the same cell. Both matrices contain all cells, *v* ∈ *𝒱*^(SG)^, where 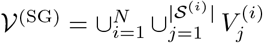 denotes the union of cells over subgraphs in all *N* tissues.

For the sCCA model associated with each *k*-hop neighborhood, *k* ∈ [1, 2, 3], we construct an input data matrix, *X*^[*k*]^, and a CNE matrix, *C*^[*k*]^. The matrices *C*^[*k*]^ and *X*^[*k*]^ are constructed by pairing replicates of each cell’s CNE with the input omics profiles of all of the cell’s neighbors at graph distance *k*. Explicitly, consider the indexed list of all pairs,

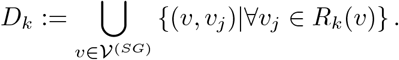

Namely, *D*_*k*_ is the indexed list of all pairs of each cell with each of the cell’s distinct neighbors at graph distance k. We construct a data matrix, 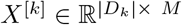, and a CNE matrix, 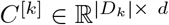, where *M* and *d* are the dimensions of the input data and CNE spaces, respectively. For the *l*-th pair, *D*_*k*_[*l*],

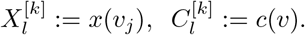

As above, *x*(*v*_*j*_) corresponds to the input cell profile of cell *v*_*j*_ and *c*(*v*) corresponds to the CNE of cell *v*. This pairing procedure generates four pairs of matrices, (*X*^[*k*]^, *C*^[*k*]^), on the input data and CNE spaces, respectively, for *k* ∈ [0, 1, 2, 3].

Sparse CCA applied to each input data-CNE matrix pair, (*X*^[*k*]^, *C*^[*k*]^) enables us to identify markers most correlated with predicted phenotype. Specifically, we adapt an *ℓ*0-based sparse CCA method to our problem[25]. This method has two hyperparameters, *λ*_*X*_ and *λ*_*C*_, associated with the input data, *X* and, CNE, *C*, matrices, respectively. The sparsity regularization controls the number of features identified in each space. We apply sparsity regularization solely to the input dimensions (*i*.*e*., *λ*_*C*_ = 0). An appropriate sparsity parameter is chosen using two-fold cross validation. The original implementation identifies a single component (the leading component). We add an orthogonal regularizer, Σ_*i*,*j*; *i*≠*j*_ |*x*_*i*_*· x*_*j*_|, where *x*_*i*_, *x*_*j*_ are canonical weight vectors, to the cost function to identify more than one component. Five components are computed for each sparse CCA model.

A local Moran’s I was computed for each marker. For a cell, *v*_*i*_, with marker expression, *x*_*i*_, the statistic, *I*_*i*_ is computed

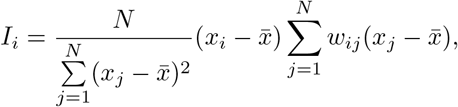

where *N* denotes the total number of cells, 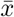 the mean expression of marker *x* across all cells, and *w*_*ij*_ is a distance-based weight. Note, the first fraction is simply the marker’s variance. Here, we let *w*_*ij*_ = 1 if *v*_*j*_ is in the three hop neighborhood of *v*_*i*_, *v*_*j*_ ∈ 𝒩_3_(*v*_*i*_). Otherwise, *w*_*ij*_ = 0. Statistics are computed sseparately for each tissue sample (*e*.*g*., 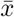 is the mean of the marker within a tissue).

To assess the significance of each statistic, we randomly permute marker expression across nodes, while retaining graphical structure. This permutation generates a distribution of permuted statistics. We repeat ten times and calculate a p value, which corresponds to the fraction of permuted statistics larger or smaller than the true statistic (the minimum of the two). As above, statistics are computed separately for each tissue. P-values were adjusted for each marker across tissues using the Benjamini-Yekutieli procedure.

### 8.8 Datasets

Here, we describe the three different disease datasets presented in the Results section: Metastatic Melanoma, Non-Small Cell Lung Cancer (NSCLC), and Colorectal Cancer (CRC). Each dataset corresponds to a specific classification task. These datasets are derived using three leading spatial transcriptomics and spatial proteomics technologies: CosMx Spatial Molecular Imager (SMI), Imaging Mass Cytometry (IMC), and Co-detection by Indexing (CODEX). Subsequent sub-sections provide details of these datasets.

#### 8.8.1 Metastatic Melanoma / CosMx Spatial Transcriptomics Classification

In this study, we generated a CosMx spatial transcriptomics dataset for the first classification task of predicting patient response to immunotherapy using metastatic melanoma samples. We initially collected 50 formalin-fixed paraffin-embedded (FFPE) tumor samples from a larger tumor microarray (TMA), representing 49 individuals (with one individual contributing two samples). Quality control measures led to the exclusion of two samples due to suboptimal data quality, and the removal of one duplicate sample. As a result, the final dataset includes 47 unique samples from 47 distinct patients.

These patients underwent treatment with one of three immunotherapy regimens: ipilimumab-nivolumab (n = 21), pembrolizumab (n = 16), or nivolumab monotherapy (n = 10). We defined responders (n = 20) as those exhibiting either a partial or complete response, as determined by comprehensive body scans. The median progression-free survival time for responders was 426.5 days (95% CI: 336, 796; estimated via 1,000 Bootstrap Samples), in contrast to 81.0 days (95% CI: 66, 136) for non-responders. These findings are visually represented in **Supplementary Figure ??a** and detailed data characteristics can be found in **Supplementary Table ??**.

Samples were sequenced using the CosMx Spatial Molecular Imager (“CosMx”) platform. CosMx sequences FFPE tissue samples via cyclic fluorescence *in situ* hybridization (FISH) techniques that iteratively identify transcripts and their location. For this dataset, the platform sequenced 960 targeted transcripts. In addition, the platform is capable of identifying a limited number of protein analytes via fluorescent immunohistochemistry; here, four cellular markers, *S100B, CD3, CD45*, and *DAPI*, were measured. The samples were sequenced over two sequencing runs (n = 26, 24, respectively). Counts per cell and negative probes identified per cell are similar between the two runs (**Supplementary Figure ??c-d**). No batch effect corrections were applied to the data.

Raw samples were processed as follows (**Supplementary Figure ??b**). First, sequencing samples were filtered to exclude cells overlapping from adjacent tumor samples in the TMA. Second, cells with fewer than ten transcripts associated with the cell were excluded. Otherwise, no additional tumor processing was conducted. Tumors were modeled as graphs via consideration of inter-cellular distance. The mean cell radius, *r*, was computed for all cells in the graph and an edge was assumed between pairs of cells within *3r* of each other. As described, two data samples were excluded for poor data quality and one sample was excluded as a (biological) replicate. The first excluded sample displayed poor sequencing depth. The second excluded sample displayed poor inter-cellular connectivity. For the biological replicate, one of the two samples was arbitrarily chosen and dropped. The samples exhibit similar sequencing quality and tissue structure. See **Supplementary Figure ??c-e** for an illustration of the excluded tissue samples.

Anchor cells were identified using the mean *S100B* protein abundance as a tumor marker and subgraphs were extracted using the described subgraph extraction algorithm (**Algorithm 1**). Cell counts, *c*_*i*_ are log-transformed, *c*_*i*_ *→* log(*c*_*i*_ + 1e− 3) and z-normalized. Per patient, an average of 9.1 (STD: 2.1) subgraphs were extracted. The model was applied to classify tissues as likely or unlikely to respond to immunotherapy. Five-fold cross validation was performed, which generated an average of 28-29, 9-10, 9-10 tissue samples for training, validation, and test sets, respectively.

#### 8.8.2 Non-small Cell Lung Cancer / Imaging Mass Cytometry Spatial Proteomics Classification

The second task, classifying response to immunotherapy of patients with non-small cell lung cancer (NSCLC) tissues, comprises a previously described dataset [70]. Protein abundances for 33 total markers were quantified using imaging mass cytometry (IMC). The data comprise 29 patients treated with PD-1 / PD-L1 immunotherapy. Patients were categorized according to response to immunotherapy as measured by full-body scans (11 responders, 18 non-responders).

Raw samples were processed as follows. Four markers, Xe-131, Xe-132, Hg-200, and 190BCKG, which represent standard background markers, were excluded. Consequently, 29 markers were used in the NSCLC analysis task. Anchor cells were identified using the pan-cytokeratin (PanCK) tumor marker and subgraphs were extracted using the described subgraph extraction algorithm (**Algorithm 1**). Cell-cell contacts, which correspond to graph edges, were pre-annotated for these data. Per patient, an average of 13.4 (STD: 5.6) subgraphs were extracted. Five-fold cross-validation was performed. On average, each fold consisted of 23, 3, and 3 tissue samples for train, test and validation, respectively. Each marker was normalized to the range of observed expression in each graph. Specifically, for a given marker *x*_*i*_, we normalize its expression levels by applying the transformation

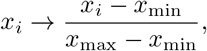

where *x*_max_ and *x*_min_ represent the observed maximum and minimum expression levels of the marker across the tissue, respectively.

#### 8.8.3 Colorectal Cancer / CODEX Spatial Proteomics Classification

The third task entails the classification of colorectal cancer (CRC) subtypes using patient-derived samples. This analysis builds upon the dataset originally generated by Schurch *et al*. [12], involving application of our proposed SORBET model to discern distinct CRC subtypes.

We do not apply any additional data processing to the raw data extracted from that work. Briefly, Schurch, *et. al*. extracted four tissue samples from FFPE samples for each patient (n = 35). Patients were categorized as displaying a Crohn’s like reaction (CLR) if two of the four tissue samples displayed tertiary lymphoid structures (TLS) and two of the four tissue samples displayed diffuse inflammatory infiltrates (DII) by hematoxylin and eosin (H&E)-stained tissue samples. Patients were categorized as displaying a DII reaction if all four tissue samples displayed DII. The CLR reaction was associated with improved survival rates when compared with the DII population. Subsequently, 56 proteins were quantified in all tissues using the CODEX technology. CODEX iteratively quantifies protein abundance via annealing of DNA-conjugated antibodies and (complementary) DNA-conjugate fluorescent probes. Raw images were processed using the CODEX Toolkit [19]. Cells were segmented based on the DRAQ5 nuclear stain using the CODEX Toolkit [12]. The dataset comprises 35 patient samples, with 17 CLR samples and 18 DII samples.

To apply SORBET, anchor cells were identified using the pan-cytokeratin (PanCK) tumor marker and subgraphs were extracted using the described subgraph extraction algorithm (**Algorithm 1**). Edges were inferred using an estimated distance threshold, *τ*. For a sample with *n* nodes, *τ* is defined as the 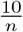-th percentile distance from the sorted list of all cell-cell distances. Cell counts, *c*_*i*_ are log-transformed, *c*_*i*_ → log(*c*_*i*_ + 1e −3), and z-normalized, as in the original paper. Per patient, an average of 35.9 (STD: 7.3) subgraphs were extracted. The model was applied to classify tissues as displaying a CLR or a DII-reaction. On average, 28, 4, and 3 tissue samples were included in the training, validation, and test sets, respectively.

