## Supplementary Figures and Tables for "SORBET: Automated cell-neighborhood analysis of spatial transcriptomics or proteomics for interpretable sample classification via GNN"

**Keywords:** spatial omics, spatial transcriptomics, graph neural networks, classification, GNN interpretability

### Appendix A Supplementary Data Figures

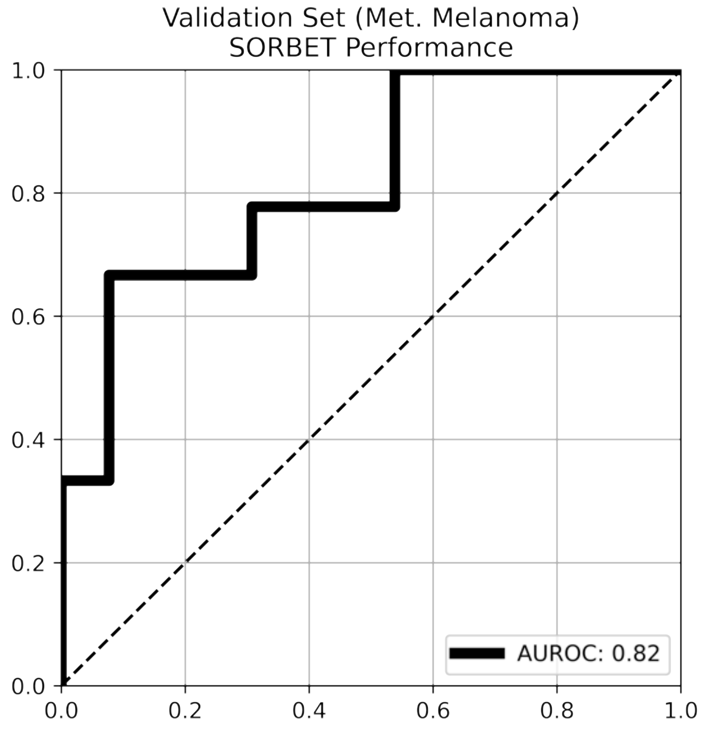


**Fig. A1 Predictive Accuracy of SORBET on Held-Out Batch for Metastatic Melanoma** Performance of SORBET predicting a held-out batch ($n = 20$ samples). Training was conducted on the first collected batch ($n = 27$ samples). Classification accuracy was assessed using a Receiver-Operator Characteristic (ROC) curve. AUROC = 0.82.

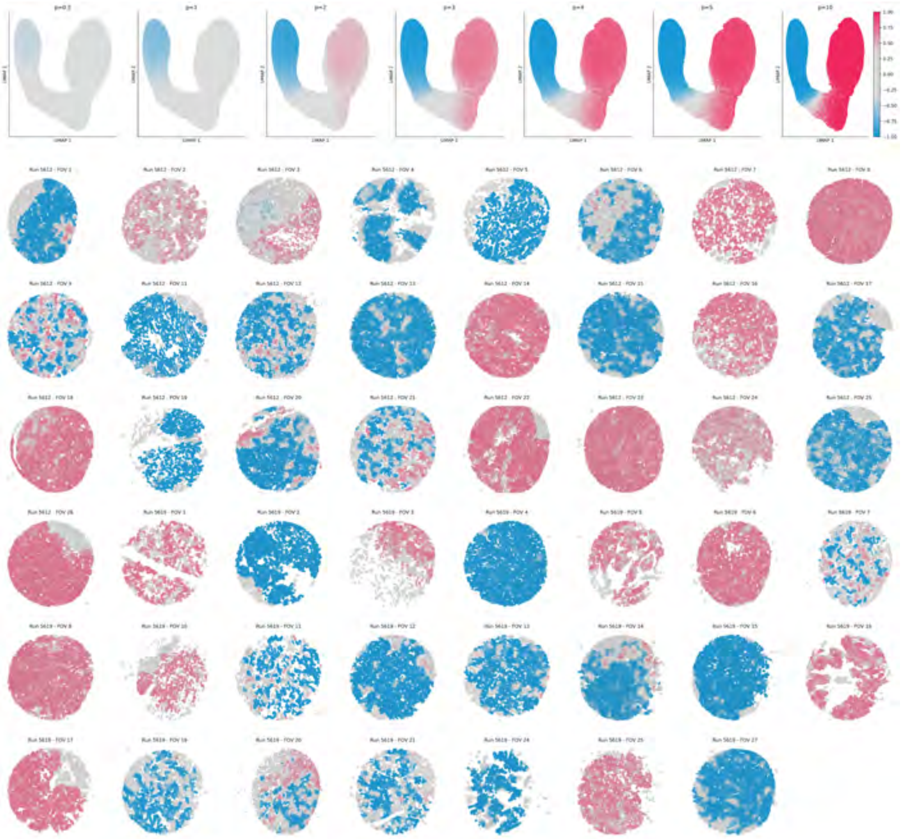

**Fig. A2 Prioritization of Spatial Patterns in Metastatic Melanoma Across all Samples** (Top) UMAP Projection of CNEs with different weight exponents $p = \{0.5, 1, 2, 3, 4, 5, 10\}$ ($p = 3$ duplicates the left column from **Figure ??**). (Bottom) Plots for all tissue samples across the dataset. The cell associated with each CNE is colored using the Cell Phenotypes Similarity score (CPS; $p = 3$). Colorscale corresponds to observed phenotype (blue - non-response to immunotherapy, grey - no association, red - response to immunotherapy).

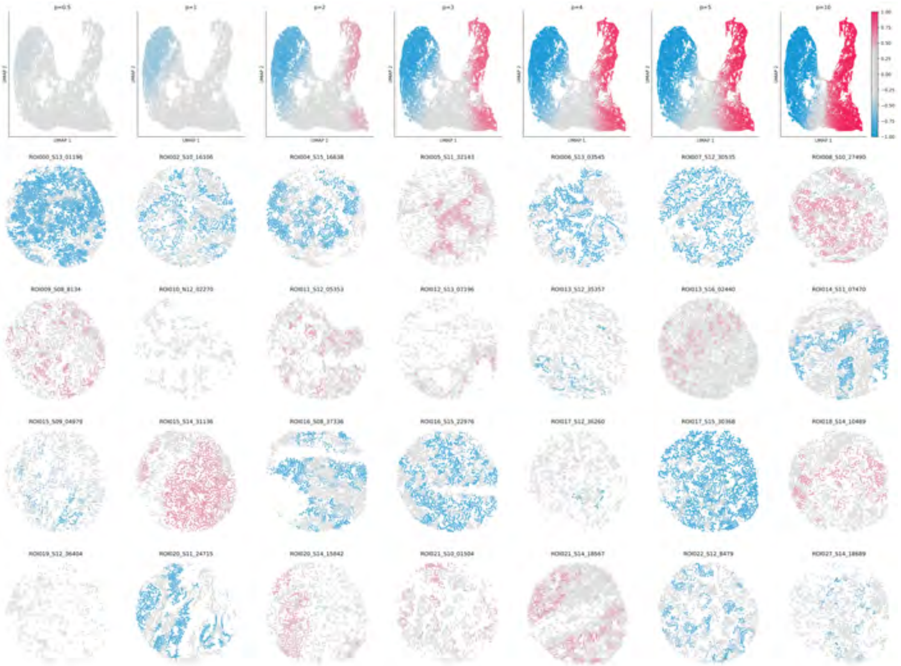

**Fig. A3 Prioritization of Spatial Patterns in Non-Small Cell Lung Cancer (NSCLC) Across all Samples** (Top) UMAP Projection of CNEs with different weight exponents $p = \{0.5, 1, 2, 3, 4, 5, 10\}$ ($p = 3$ duplicates the left column from **Figure ??**). (Bottom) Plots for all tissue samples across the dataset. The cell associated with each CNE is colored using the Cell Phenotypes Similarity score (CPS; $p = 3$). Colorscale corresponds to observed phenotype (blue - non-response to immunotherapy, grey - no association, red - response to immunotherapy).

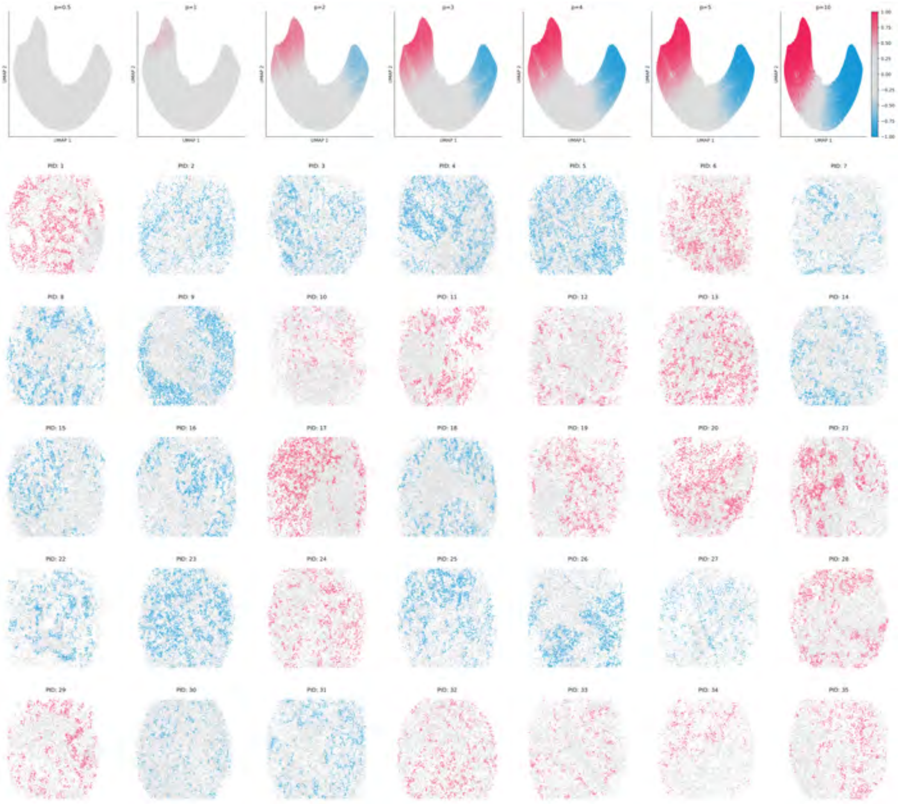


**Fig. A4 Prioritization of Spatial Patterns in Colorectal Cancer (CRC) Across all Samples** (Top) UMAP Projection of CNEs with different weight exponents $p = \{0.5, 1, 2, 3, 4, 5, 10\}$ ($p = 3$ duplicates the left column from **Figure ??**). (Bottom) Plots for all tissue samples across the dataset. The cell associated with each CNE is colored using the Cell Phenotypes Similarity score (CPS; $p = 3$). Colorscale corresponds to observed phenotype (blue - non-response to immunotherapy, grey - no association, red - response to immunotherapy).

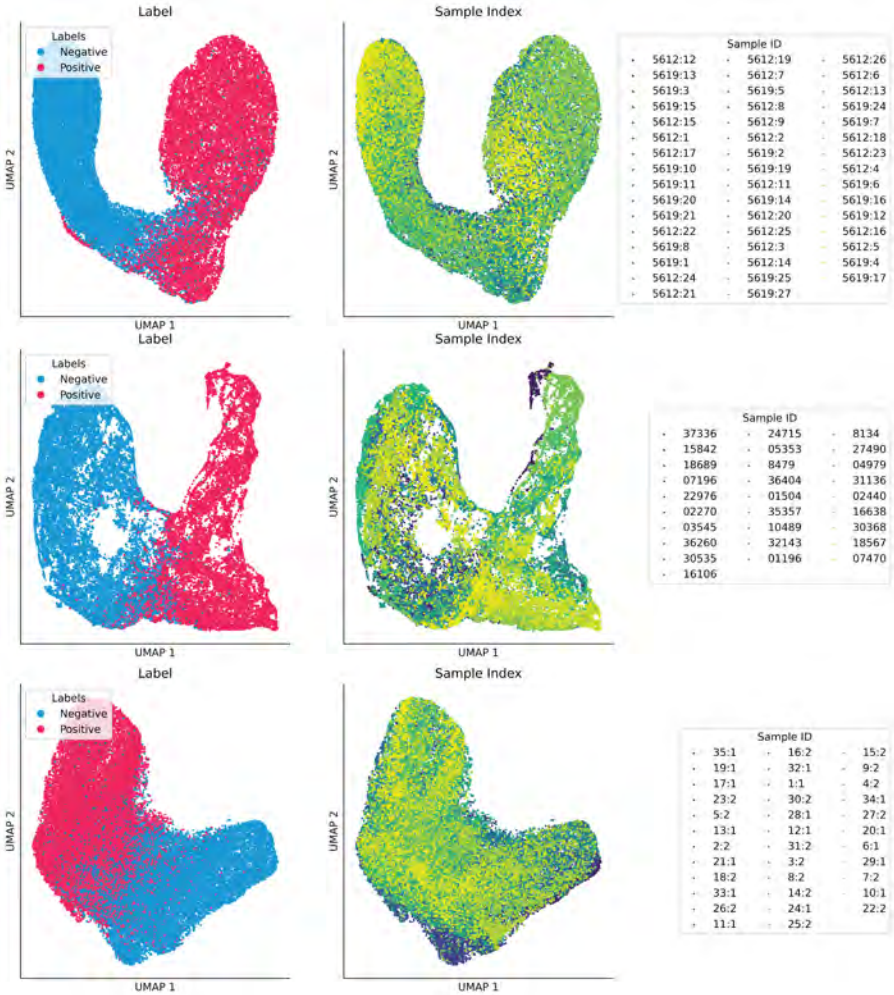


**Fig. A5 Cell Niche Embeddings: Phenotypic and Sample Labels** UMAP Projection of CNEs colored according to subgraph label (left) and tissue sample ID (right) for (top) Metastatic Melanoma, (middle) Non-Small Cell Lung cancer, and (bottom) Colorectal cancer.

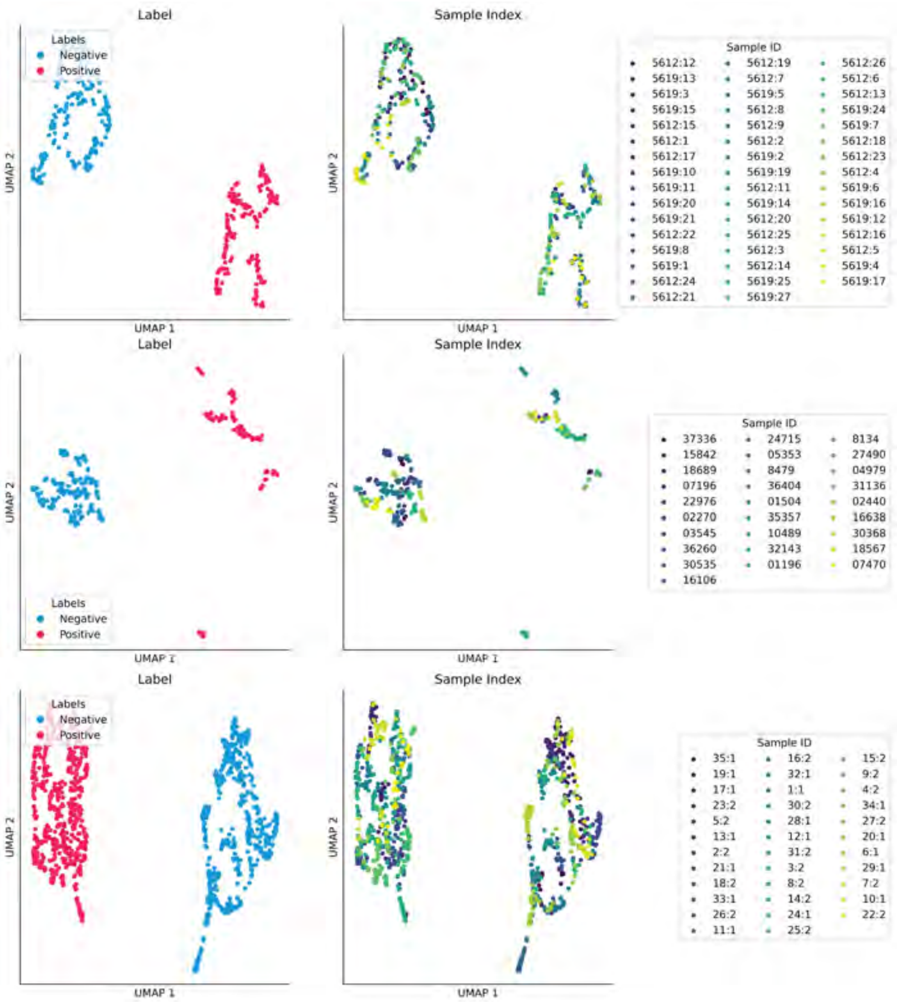


**Fig. A6 Subgraph Embeddings: Phenotypic and Sample Labels** UMAP Projection of SEs colored according to label (left) and tissue sample ID (right) for (top) Metastatic Melanoma, (middle) Non-Small Cell Lung cancer, and (bottom) Colorectal cancer.

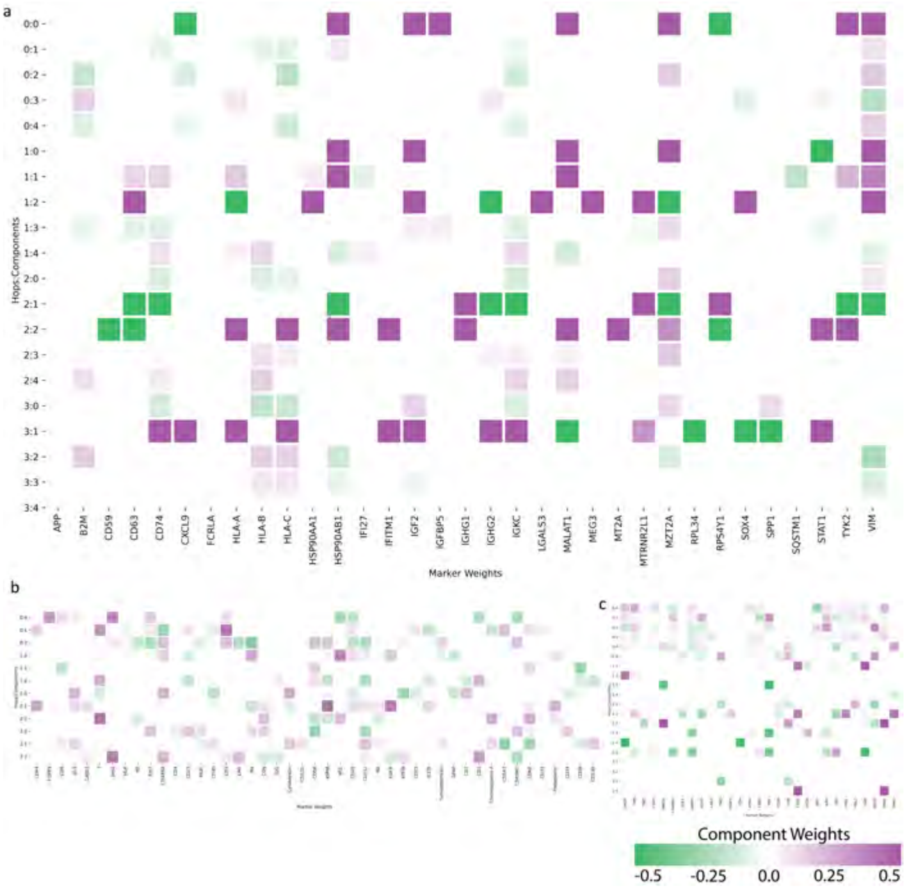


**Fig. A7 Complete Marker Programs** Complete marker programs for each of the three datasets: **(a)** Metastatic Melanoma - CosMx, **(b)** NSCLC - IMC, **(c)** CRC - CODEX. Rows define individual components. Components are annotated by hop distance and component index using the structure 'Hop Distance':'Component Index' (*e.g.,* 2:3 denotes the third computed component of the two-hop sCCA model). Columns are annotated by marker. **(a)** Includes the top 30 non-zero weight norm markers for metastatic melanoma (n = 960 total markers).

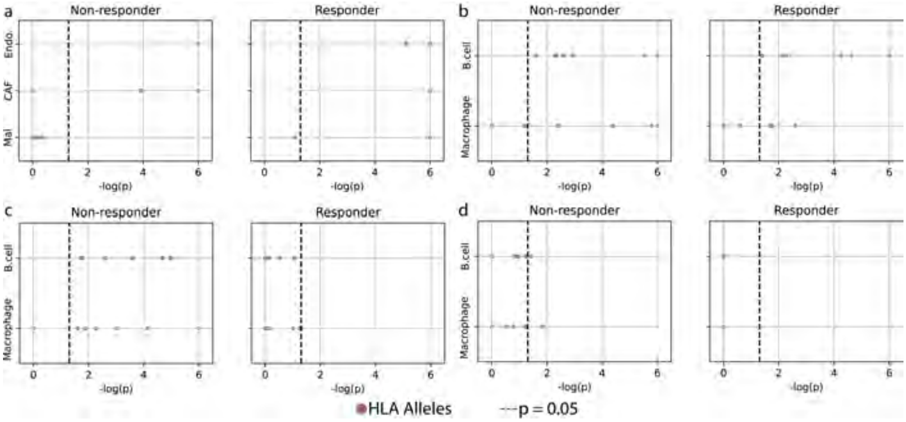


**Fig. A8 Analysis of CPS Scores in the Context of T Cell Activation** . Additional analyses of changes in CPS scores in response to activation. Corresponds to **Fig. ??f-h**. Sub-figures correspond to activation of **(a)** $CD8^+$ T cells (same as **Fig. ??g**), **(b)** Naive $CD4^+$ T cells **(c)** Th1 $CD4^+$ T cells (same as **Fig. ??h**, and **(d)** Th1 $CD4^+$ T cells. APCs are plotted on the y-axis. Adjust $-\log_{10}(p)$ value (Benjamin-Yekutieli procedure) is plotted on the Y-axis. Non-responders (left) and responders (right) are analyzed separately.

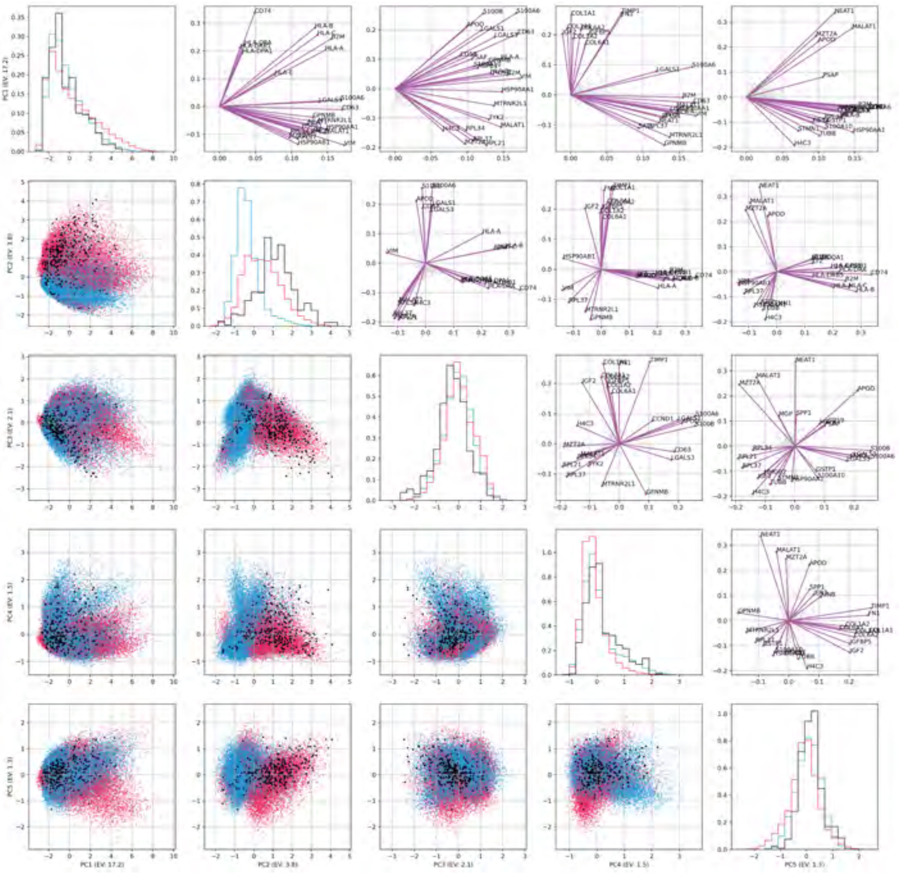

**Fig. A9 Comparison of Expression Profiles in Each Cell**. A principle component analysis (PCA) of the expression profiles of each cell. Each row and column corresponds to a principle component (leading five principle components). Plots may be split by plots below the diagonal, plots along the diagonal, and plots above the diagonal. **Below the diagonal:** Scatter plots of two principle components corresponding to the row and column. Each point is colored by the label of the source tissue (red - responder, blue - non-responder). Negatively labeled points with positive CPS scores are colored black. **Diagonal:** Histograms comparing each population (red - responder, blue - non-responder, black - non-responder with positive CPS) for each component. **Above the diagonal:** Scatter plot demonstrating the loadings of pairwise principle components. The top 20 markers (by magnitude of loadings) are highlighted by purple lines and labeled. Otherwise, each marker is represented by a grey point.

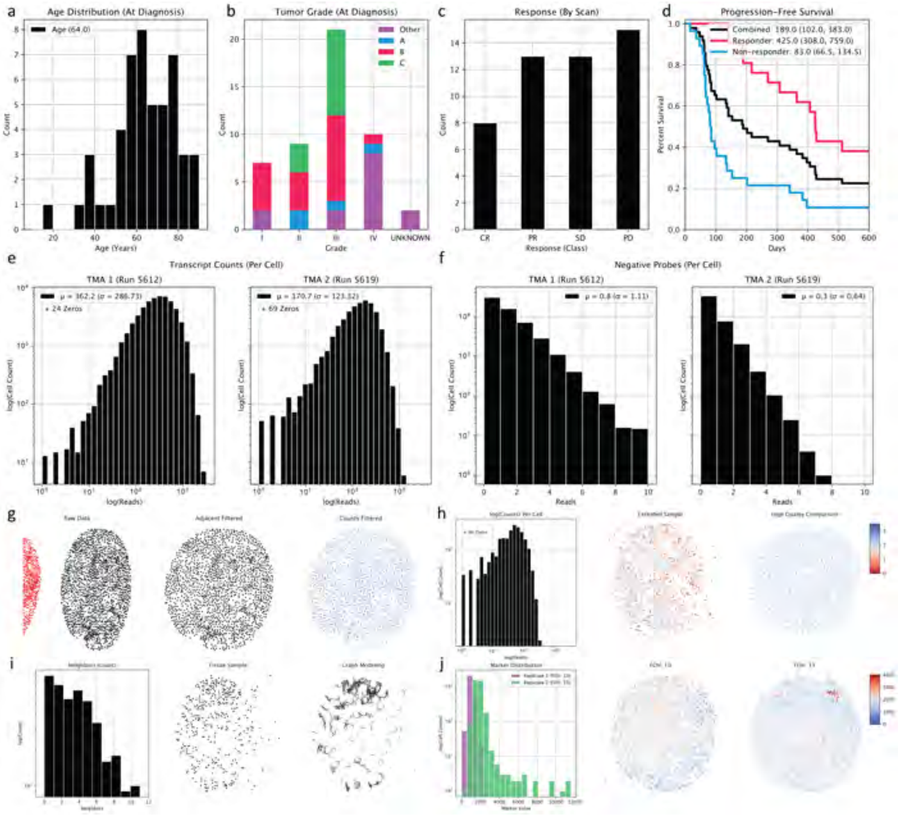


**Fig. A10 Metastatic Melanoma Data Quality (a-d)** Data-set Characteristics. **(a)** Age distribution. **(b)** Tumor grade at diagnosis. Grade sub-type is reported using the colors (*i.e.* Count of IIA is reported in blue bar for Grade II tumors). **(c)** Response class. **(d)** Kaplan-Meier Curves including Overall Progression-free Survival (PFS) and PFS among Responder and Non-responder phenotypes. Median PFS is reported in the legend with a 95% Confidence Interval computed using Bootstrap Iteration (n = 100,000 iterations). **(e)** Distribution of transcript counts per cell (log-log) for the two sequencing runs. **(f)** Distribution of transcript counts per cell (count-log) for the negative sample probes. **(g)** Overview of tissue processing steps: Overlapping sample boundaries (red) are removed from the 'Raw Data' manually. Cells with low transcript counts are (subsequently) removed from the sample. **(h)** Tissue excluded for low-sequencing quality. (left) Transcript counts per cell (log-log) for the tissue (analogous plot to **(e)** on the sample's subset of cells). Sample plots colored according to transcript counts for (middle) poor quality / excluded sample and (right) higher-quality sample. **(i)** Tissue sample removed from analysis due to poor graph modeling. Distribution of edge weights with appropriate threshold (left) with raw cell layout (middle) and graph-modeled tissue (right) displayed. **(j)** Illustration of Mean *S100B* value measured for for duplicated sample. (left) Distribution of Mean *S100B* ('Marker Value') for both samples. (middle, right) Sample plots of duplicated sample. 'FOV 15' was retained.

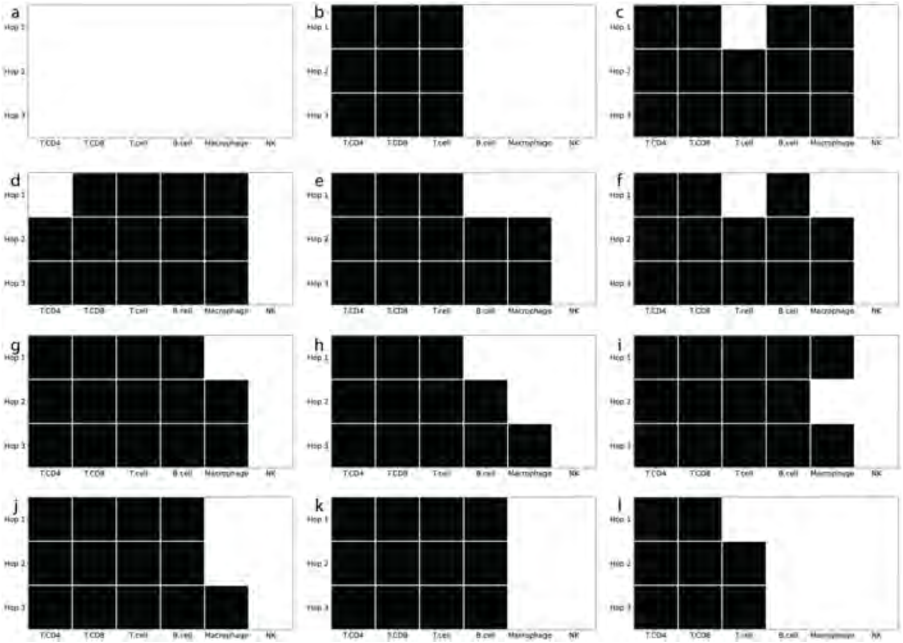


**Fig. A11 Significant TCN Patterns Identified Using CPS (a)** Binary heatmaps of significant TCN patterns identified using the CPS scores in the TCN analysis. Each pattern is associated with a TCN cluster including many cell niches. Significant cell type-distance pairs and markers are plotted in **Fig. A12** and **Fig. A13**, repectively

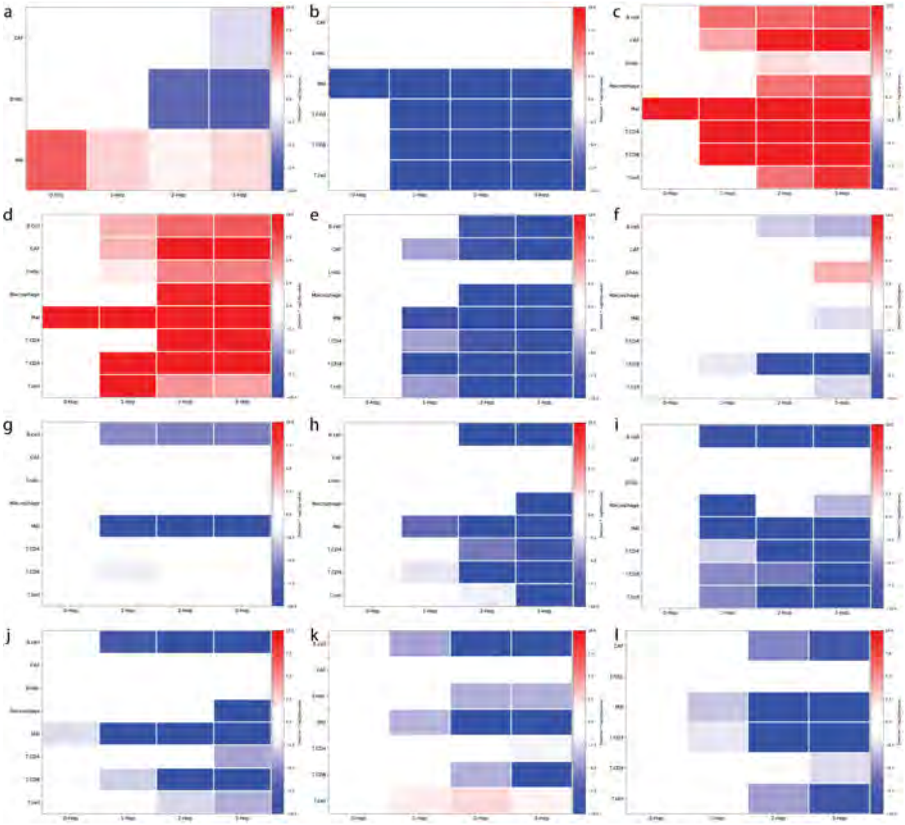


**Fig. A12 Significant Cell Type-Distance Pairs TCN Patterns Identified withing Significant TCNs** Heatmap of significant CT-Distance pairs for the TCN pattern in **Fig. A11**, with blue representing significance towards non-responders and red indicating significance towards responders. The magnitude corresponds to $-\log_{10}$(p-value).

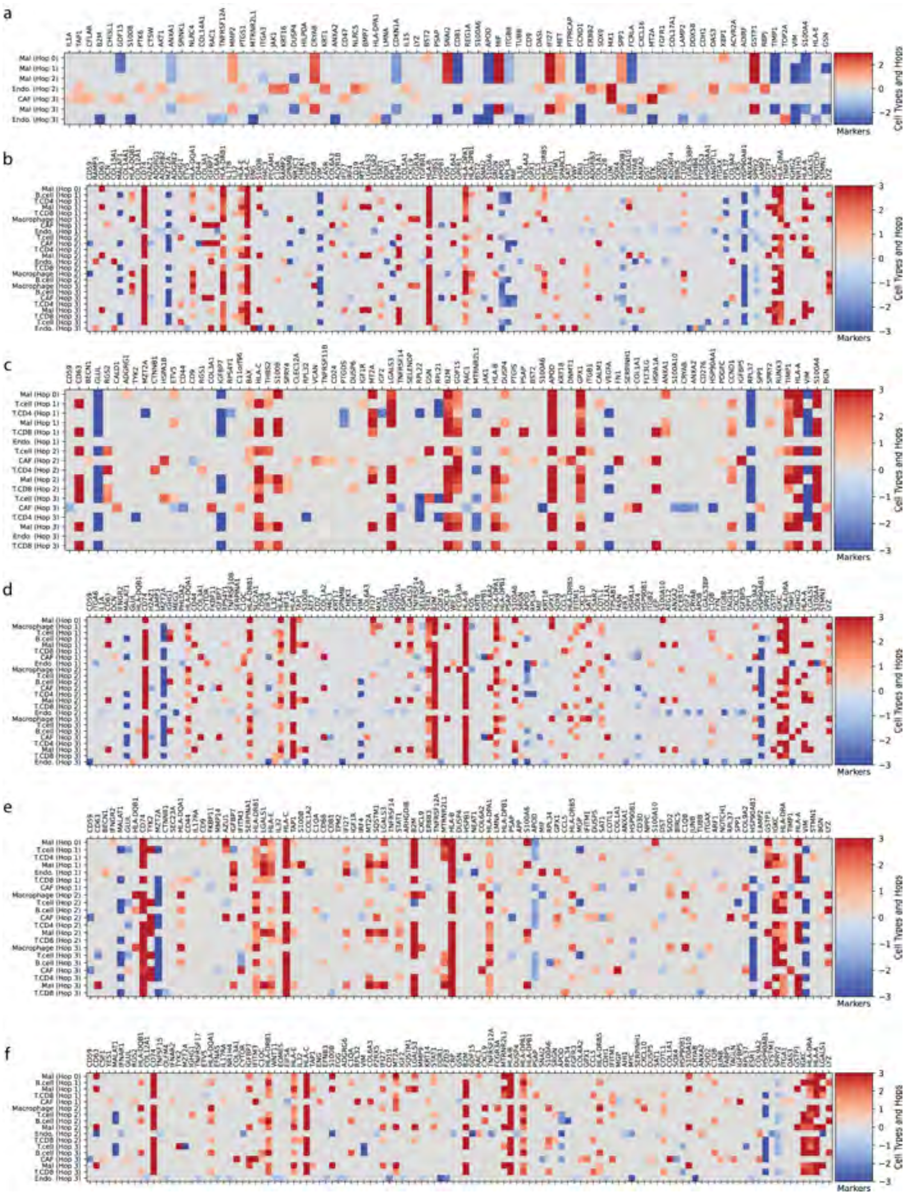


**Fig. A13 Differentially expressed markers for each significant cell type in each significant TCN.** Heatmap of significant markers for the TCN clusters associated with the patterns in **Fig. A11**. Rows represent significant CT-Distance pairs (from **Fig. A12**), and columns correspond to markers. Similarly, red and blue correspond to significance towards responders and non-responders, respectively. The magnitude corresponds to mean expression value. Subplots share the same indexing with **Figures A11, A12** (*e.g.*, **(a)** in this figure corresponds to **(a)** in those figures). Continued in **Fig. A14**

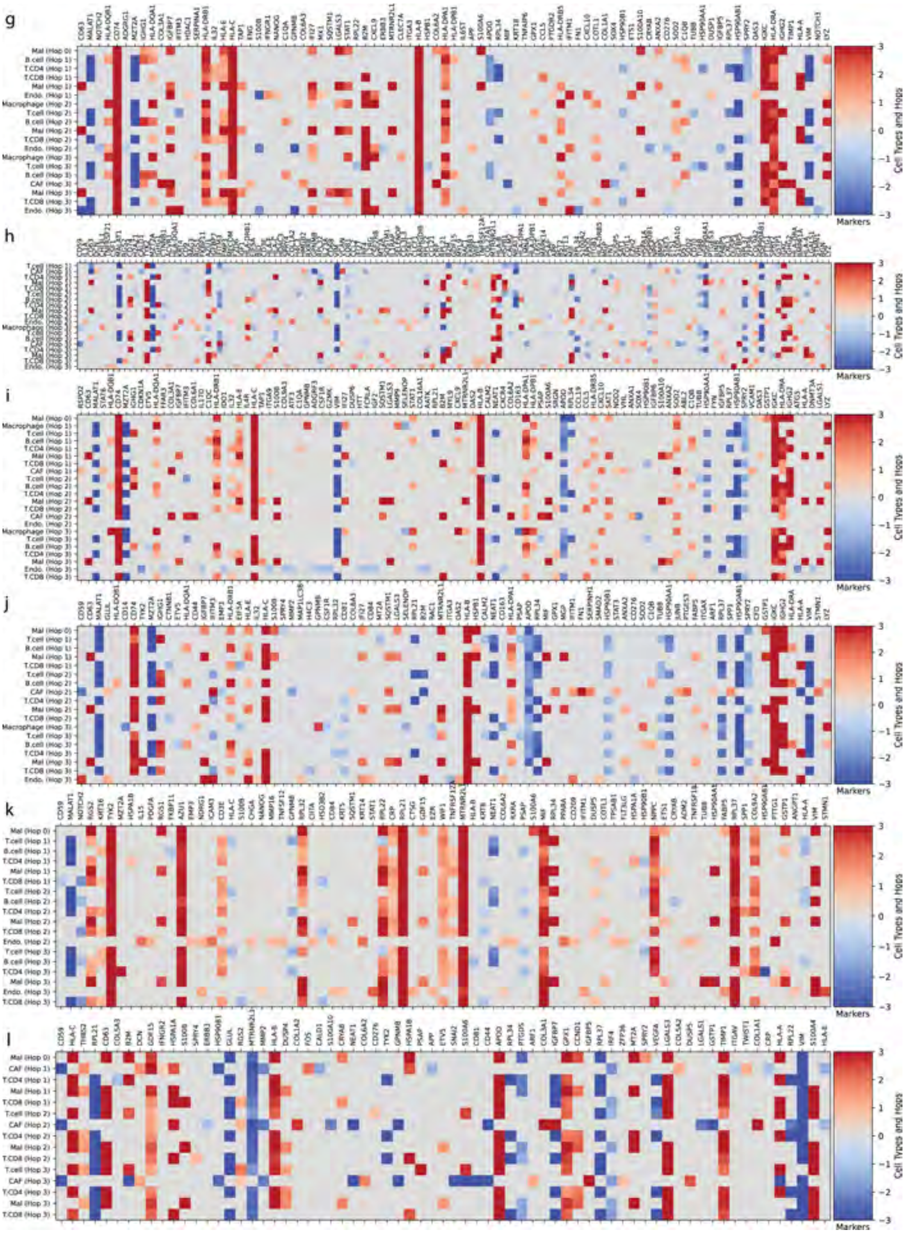


**Fig. A14 Differentially expressed markers for each significant cell type in each significant TCN (Continued).** Continuation of Fig. A14. See that figure caption for details.

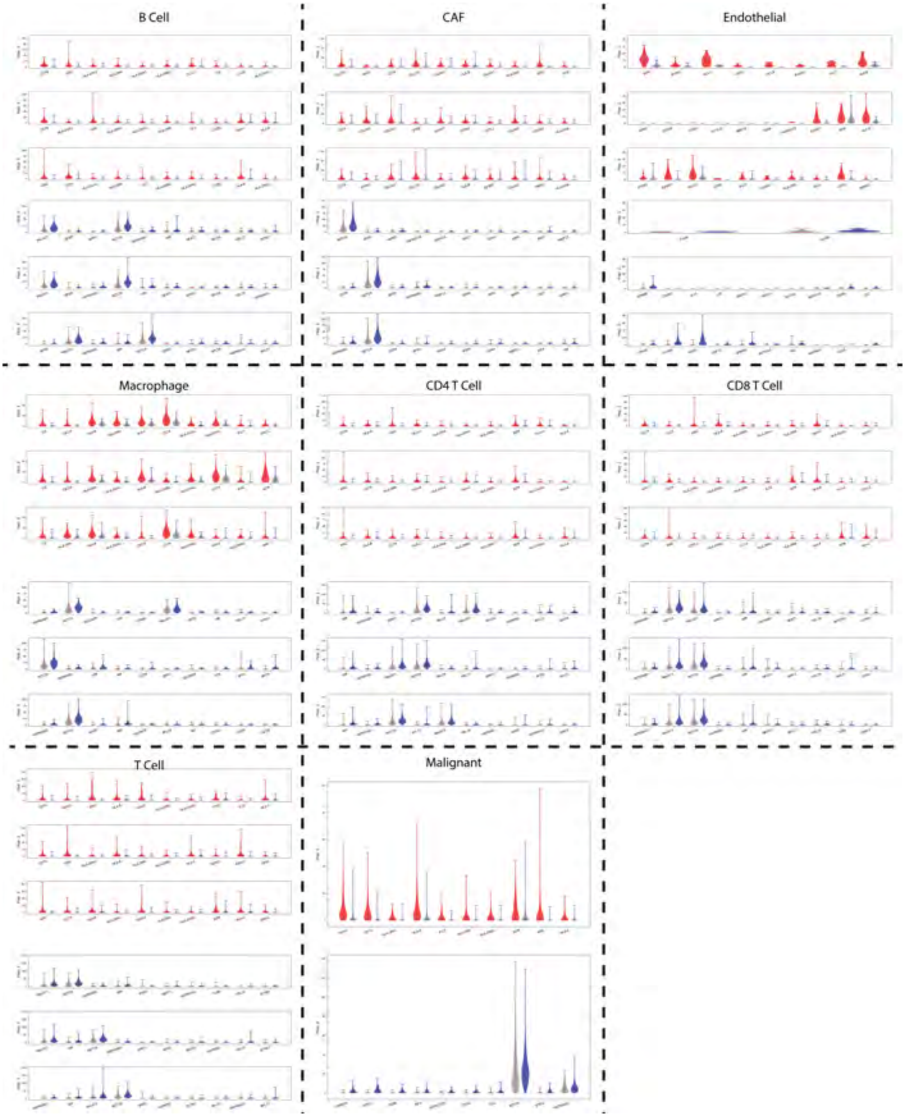


**Fig. A15 Top Ten Differentially Expressed Markers per Cell Type.** Violin plots of the expression levels for the ten most differentially expressed markers across all TCNs in each cell type. Cell types are separated by dashed lines with annotations at the top of the box. Significant marker sets are identified for responders (red, top) and non-responders (blue, bottom). In each plot, marker expression in significant cell niches with the opposite label is plotted in grey (*e.g.*, Non-responders in the top plot). Corresponds to **Figure ??i,j**.

### Appendix B Supplementary Data Tables

| | **Responder** | **Non-Responder** |
|---|---|---|
| **Total (N=47):** | 20 | 27 |
| **Sex (% Male):** | 65 | 56 |
| **Mean Age (STD):** | 62.7 (13.7) | 64.6 (17.2) |
| **Stage At Diagnosis** | | |
| I | 5 | 2 |
| II | 5 | 3 |
| III | 6 | 14 |
| IV | 3 | 7 |
| Unknown | 1 | 1 |
| **Best Response** | | |
| Complete Response | 8 | 0 |
| Partial Response | 12 | 0 |
| Stable Disease | 0 | 12 |
| Progressive Disease | 0 | 15 |
| **Therapy** | | |
| Ipi / Nivo | 8 | 13 |
| Pembrolizumab | 8 | 8 |
| Nivolumab | 4 | 6 |
| **Prior Checkpoint Blockade** | | |
| Yes | 6 | 6 |
| No | 14 | 21 |
| **Annotated Mutation** | | |
| BRAF | 6 | 7 |
| *V600E* | 5 | 6 |
| *V600K* | 1 | 1 |
| NRAS | 2 | 5 |
| *Q61R* | 2 | 0 |
| *Q61K* | 0 | 1 |
| *Q61L* | 0 | 1 |
| *Q61H* | 0 | 1 |

**Table B1 Summary of the characteristics of the Melanoma dataset.** Data are split into "Responder" and "Non-responder" phenotypes and Demographic information (Sex, Age), Stage at Diagnosis, Best Response, Therapy, History of Checkpoint Blockade, and Mutation Status are highlighted.

| | NSCLC | CRC | Melanoma |
|---|---|---|---|
| #Patients | 29 | 35 | 47 |
| #Tissues | 29 | 140 | 47 |
| #Features | 29 | 49 | 960 |
| #Positive/#All | 11/29 | 17/35 | 20/47 |
| Classification Task | PD-1 R. | Tissue Phenotype | ITx R. |
| Type | Proteomics | Proteomics | Transcriptomics |
| Technology | IMC | CODEX | CosMx |

**Table B2 Summary of the datasets comprising the three prediction tasks.** In classification tasks, 'R.' denotes response to therapy and 'ITx' denotes immunotherapy. Technologies are Imaging Mass Cytometry (IMC), CO-Detection by indEXing (CODEX) and the CosMx Spatial Molecular Imager ('CosMx') platform.

| Module | Parameter | NSCLC | CRC | Melanoma |
|---|---|---|---|---|
| **Embedding** | Layers | 1-3 | 1-3 | 1-3 |
| | Width | 8-32 | 16-48 | 48-96 |
| **GCN** | Layers | 2-5 | 2-5 | 2-5 |
| | Width | 16-32 | 16-48 | 48-96 |
| **CNE** | Layers | 1-3 | 1-3 | 1-3 |
| | Width | 32-64 | 96-192 | 96-192 |
| **Prediction** | Operator | Max | Max | Max |
| | Layers | 1 | 1 | 1 |
| | Width | 8-32 | 16-48 | 96-192 |

**Table B3 Model Structure Search Space for Each Classification Task.** Module corresponds to the module descriptions in Methods **??**. Layers denotes the number of intermediate representations in each module; width denotes the potential width of each transformation; operator denotes the non-linearity applied across all vertices in the graph. Here, we choose to take the max across all layers.

| CellType/Group | Unique CPS Markers | Unique Original Markers |
|---|---|---|
| B.cell/R | *EIF5A*, *GLUL*, *GSN*, *LGALS1*, *S100A10* | *MZT2A* |
| B.cell/NR | *CRYAB*, *DUSP6*, *GPX1*, *NOTCH2* | *AZU1*, *CALM2*, *CD81*, *H4C3*, *TYK2* |
| Mal/NR | *COL18A1*, *NR1H2*, *EIF5A*, *MIF*, *MT2A*, *RGS1*, *BMP7*, *LGALS3*, *PSAP*, *CALD1*, *MGP*, *C9orf16*, *G6PD*, *IER3*, *EZR*, *HSPB1*, *HSP90AB1*, *HSPA1A*, *S100A10*, *S100B*, *LGALS1*, *AZU1*, *RPL22*, *S100A6*, *DUSP6*, *CRYAB*, *KIT* | *HLA-B*, *APP*, *HLA-E*, *ITGA3*, *ITGA6*, *CD276*, *IL16*, *ADGRG1*, *GDF15*, *MMP14*, *TNFRSF10B*, *S100A4*, *ERBB3*, *ITGB8* |
| Mal/R | *CSK*, *IGF2R*, *IGHG1*, *JAK1*, *MET*, *MMP2* | *C9orf16*, *IL6R*, *KRT14* |
| T.CD8/NR | *CD44*, *CRYAB*, *DUSP4*, *DUSP6*, *HSD3B2*, *KRT18*, *LAMP2*, *SOX4* | *ADGRG1*, *CCND1*, *GSN*, *JUNB*, *LGALS1*, *S100A6* |
| T.CD8/R | *ANXA2*, *RGS1* | *ARHGDIB*, *HSPB1*, *IGF2R*, *IL6ST*, *NEAT1* |
| T.cell/NR | *GPNMB*, *IRF4*, *MGP*, *PTGDS*, *SPP1* | *CRP*, *EIF5A*, *GPX1*, *NDRG1*, *TIMP1* |
| T.cell/R | *EIF5A*, *HLA-DPB1*, *LYZ*, *VIM* | *APP*, *IL6ST*, *MZT2A*, *SPRY2* |
| T.CD4/NR | *GPNMB*, *HLA-B*, *KRT8* | *B2M*, *GSN*, *H2AZ1*, *HLA-E*, *HSPB1*, *IFITM1*, *LMNA*, *TIMP1* |
| T.CD4/R | *CALM2*, *CYTOR*, *ETV5*, *HSP90AA1*, *IFI27*, *RAC1* | *GPNMB*, *LGALS3BP* |
| Macrophage/NR | *APP*, *AZU1*, *CD14*, *CSF1R*, *ETV5*, *RAC1*, *SELENOP*, *SPRY2*, *STMN1* | *ANXA1*, *BST2*, *BTG1*, *CD44*, *CD81*, *FABP5*, *HSP90AA1*, *HSPB1*, *IFITM1*, *LGALS1*, *PSAP*, *S100A6*, *SQSTM1*, *TIMP1*, *TUBB*, *TYK2* |
| Macrophage/R | *AZU1*, *GSTP1*, *RGS1*, *SRGN* | *CD14*, *CD163*, *CD63*, *FYB1*, *JUNB*, *PTGDS*, *RAC1*, *ZFP36* |
| CAF/NR | *CD68*, *ERBB3*, *LGALS1* | *AZU1*, *CCND1*, *CD9*, *COL1A1*, *COL1A2*, *COL3A1*, *CRYAB*, *GSN*, *HLA-C*, *IGFBP7*, *ITGB1*, *TIMP1* |
| CAF/R | *ANXA2*, *BMP7*, *CHI3L1*, *COL14A1*, *CTSW*, *CXCL16*, *DDX58*, *DUSP1*, *ENG*, *IL17RA*, *IL1A*, *LYZ*, *MET*, *MX1*, *NLRC4*, *NOTCH1*, *PPARG*, *PTPRCAP*, *RARRES2*, *REG1A*, *SEC23A*, *SPARCL1*, *SPINK1*, *TAP1* | *ACTG2*, *ADGRF3*, *APOD*, *ATG5*, *ATM*, *ATR*, *AXL*, *BAX*, *BCL2*, *BTG1*, *C11orf96*, *C1QA*, *CCL2*, *CD164*, *CD55*, *CD58*, *CD59*, *CD83*, *CELSR1*, *CLEC2B*, *CLU*, *COL5A2*, *COL8A1*, *CXCL3*, *DUSP4*, *EPHA4*, *EPHB3*, *EPOR*, *ERBB3*, *ETV5*, *FGF9*, *FKBP11*, *FYN*, *G6PD*, *GLUD1*, *GZMK*, *HLA-A*, *HLA-DQB1*, *IER3*, *IFNGR1*, *IGF1*, *IGF1R*, *IL10RB*, *IL11RA*, *IL13RA1*, *IL16*, *IL1R1*, *IL1RAP*, *IL20RA*, *IRF4*, *ITGAL*, *ITGB5*, *ITGB8*, *KRAS*, *KRT8*, *LAIR1*, *LAMP2*, *LAMP3*, *LINC02446*, *MKI67*, *NDRG1*, *NLRP2*, *NLRP3*, *PDGFC*, *PIGR*, *PPARA*, *PRSS2*, *RXRB*, *S100B*, *SELENOP*, *SMAD4*, *SOD1*, *ST6GAL1*, *ST6GALNAC3*, *TLR8*, *TNFRSF10B*, *TOP2A*, *VEGFA*, *VEGFB* |
| Endo/NR | *ACE*, *ADGRA3*, *ADGRD1*, *ADGRF4*, *ADGRG6*, *AHI1*, *ANGPT1*, *ANXA4*, *CALM3*, *CCL28*, *CD19*, *CD24*, *CD47*, *CD55*, *CD79A*, *COL12A1*, *COL14A1*, *COL8A1*, *CSF3*, *CYSTM1*, *EFNA5*, *EFNB1*, *EFNB3*, *EOMES*, *ERBB2*, *ESR1*, *ETV4*, *FASLG*, *FGF1*, *FGFR3*, *FGG*, *FZD3*, *GPER1*, *HCST*, *IFNAR1*, *IGF2*, *IL10*, *IL10RB*, *IL16*, *IL17A*, *IL17B*, *IL2RG*, *IRF4*, *ITGB4*, *KLRK1*, *KRT23*, *LY6D*, *MMP9*, *MYH11*, *NGFR*, *NLRP2*, *NPR2*, *NR1H4*, *NR3C1*, *OLFM4*, *P2RX5*, *PDGFA*, *PHLDA2*, *PIGR*, *PTGES*, *PTTG1*, *RARG*, *RNF43*, *SMAD4*, *SMO*, *SOX2*, *STAT5B*, *TGFB2*, *TGFBR1*, *TLR1*, *TNFRSF11B*, *TNFRSF17*, *TNFRSF19*, *TTR*, *WNT11* | *ANXA1*, *AZU1*, *B2M*, *BCL2L1*, *BST2*, *BTG1*, *C11orf96*, *CALD1*, *CALM2*, *CD164*, *CD58*, *CD81*, *COL4A1*, *COTL1*, *CTNNB1*, *CXCL12*, *CXCL2*, *DUSP4*, *EIF5A*, *EMP3*, *FAS*, *FOS*, *FZD1*, *GPX1*, *GSN*, *GSTP1*, *H2AZ1*, *HLA-A*, *HLA-B*, *HSP90AA1*, *HSP90AB1*, *HSPB1*, *ITGA2*, *ITGB1*, *LGALS1*, *MIF*, *MMP14*, *MTOR*, *MZT2A*, *NDRG1*, *NR1H2*, *RPL21*, *RPL32*, *S100A10*, *S100A4*, *SEC61G*, *SNAI2*, *SOD1*, *SOD2*, *SPRY2*, *SQSTM1*, *TNFRSF14*, *TPM1*, *TSC22D1*, *TUBB*, *TYK2*, *UBE2C* |
| Endo/R | *AKT1*, *APP*, *COL17A1*, *DCN*, *EFNB2*, *EMP3*, *ERBB2*, *FGFR1*, *GPX1*, *IFNGR1*, *JAK2*, *KRT1*, *KRT16*, *LGALS1*, *NLRC5*, *OASL*, *PTGS1*, *PTK6*, *RBPJ*, *SOX9*, *TAP1*, *THBS2*, *TOP2A* | *ARF1*, *ARHGDIB*, *BAG3*, *BGN*, *BST2*, *C11orf96*, *C1QA*, *CCL11*, *CCL21*, *CCL3L3*, *CDH5*, *COL1A1*, *COL3A1*, *COL5A2*, *COL6A3*, *CSF3*, *CXCL10*, *FLT1*, *FPR1*, *GSTP1*, *HIF1A*, *HLA-DQA1*, *HMGB2*, *IGFBP5*, *ITGAE*, *JUNB*, *KDR*, *KLF2*, *LYZ*, *MALAT1*, *MGP*, *MMP2*, *MYL9*, *NOTCH1*, *NOTCH3*, *PTGDR2*, *PTGES3*, *RGS5*, *SLC40A1*, *SPRY4*, *SRGN*, *TAGLN*, *TNFAIP6*, *TNFRSF1B*, *TNFRSF21*, *TP53*, *TPM1*, *TYROBP*, *WIF1*, *ZFP36* |

**Table B4** Unique markers for CPS and Original by Cell Type and Group (R=Responder, NR=Non-Responder)

### References
